# Metabolic and Population Profiles of Active Subseafloor Autotrophs in Young Oceanic Crust at Deep-Sea Hydrothermal Vents

**DOI:** 10.1101/2025.04.25.650746

**Authors:** Sabrina M. Elkassas, Caroline S. Fortunato, Sharon L. Grim, David A. Butterfield, James F. Holden, Joseph J. Vallino, Christopher K. Algar, Lisa Zeigler Allen, Benjamin T. Larson, Giora Proskurowsk, Emily Reddington, Lucy C. Stewart, Begum Topçuoğlu, Julie A. Huber

## Abstract

At deep-sea hydrothermal vents, magmatically driven rock-water reactions in the crust generate gases and other reduced compounds that subseafloor microorganisms use for chemolithoautotrophy. In this study, microbial autotrophs from three diffuse flow hydrothermal vents at Axial Seamount in 2013 and 2014 were isotopically labeled using RNA Stable Isotope Probing (RNA-SIP), targeting subseafloor autotrophic mesophiles (30°C), thermophiles (55°C), and hyperthermophiles (80°C). We constructed taxonomic and functional profiles of active chemolithoautotrophs, examined population distributions across sites, and linked primary producers to their specific metabolic strategies within the subseafloor community. Dominant autotrophs exhibited hydrogen-dependent dissimilatory metabolisms such as sulfur and nitrate reduction and methanogenesis, as well as microaerophilic sulfide oxidation even at 80°C, consistent with fluid chemistries at each site. While hydrogenotrophic methanogenic archaea such as *Methanothermococcus* were restricted in their distribution and activity, hydrogenotrophic sulfur and nitrate reducers from the Aquificota (*Thermovibrio*), Campylobacterota (Nautiliaceae, *Hydrogenimonas*, and Desulfurobacteriaceae) were consistently active and present at all sites and years at both the population and community levels. Hydrogenase transcripts were significantly differentially expressed, and diverse hydrogenases were found in metagenome-assembled genomes of Aquificota members, highlighting the importance and versatility of their hydrogen utilization strategies which likely contribute to their cosmopolitan distribution across geochemically disparate subseafloor sites. Together, this study provides new insights into the functional dynamics and distribution of key subseafloor autotrophic microbial communities in young oceanic crust at deep-sea hydrothermal vents.

**IMPORTANCE:** Deep-sea hydrothermal vents are hotspots for life in the dark ocean, where rich animal ecosystems are supported by microbial primary producers utilizing the abundant chemical energy supplied by high-temperature water-rock reactions. Despite increasing knowledge about the geochemistry and microbiology of deep-sea hydrothermal vents, there is still a gap in our understanding of the key microbial players who fix much of the carbon at these sites, especially in the productive subseafloor. In this study, stable isotope probing was used to label active microbial autotrophs in diffuse flow venting fluids from three sites over two years and was combined with metatranscriptomic sequencing to identify their specific metabolic strategies. This research highlights the microbial community composition, function, gene regulation, and population dynamics that enable hydrothermal ecosystems to persist.

## INTRODUCTION

Deep-sea hydrothermal systems span large temperature and geochemical gradients, from cold, oxidized seawater to hot, highly reduced hydrothermal fluid produced by rock-water reactions. These systems host a wide diversity of microbial niches, many of which are powered by chemolithoautotrophy, including chimneys, animals, venting fluids, hydrothermal plumes, mats, and the subseafloor (1). Mixing zones at and below the seafloor can be sampled through diffuse flow vents (∼5-120°C), providing a glimpse into the subseafloor habitat within oceanic crust, one of the most productive microbial niches in the vent environment (2–11).

Phylogenetic, metagenomic, and metatranscriptomic studies of diffuse flow microbes have expanded our knowledge on the composition and variability of these communities over time and space, spanning a variety of tectonic settings, host rock types, and geochemical regimes (e.g. 1, 7-10, 12-22). The diffuse flow microbial communities identified in these studies include a mix of seawater, seafloor, and subseafloor populations that inhabit a wide range of temperatures along diverse geochemical gradients, making it difficult to identify active subseafloor residents and the factors controlling their distribution and metabolisms. However, the recent application of stable isotope probing, including RNA Stable Isotope Probing (RNA-SIP), is providing new insights into microbial activity in the dynamic and hard-to-access subseafloor environment (8, 11, 23–24).

RNA-SIP is an experimental technique by which isotopically labeled substrate is added to environmental samples at near-natural conditions, incubated, and the label incorporation is probed at the genetic level through sequencing of transcripts (8, 25–26). Resulting isotope-enriched RNA-SIP metatranscriptomes reveal the identity and functional attributes of the microbes that incorporate the labeled substrates in each community under defined experimental conditions (8, 25–26). The first deep-sea application of this method was at Axial Seamount (Juan de Fuca Ridge NE Pacific Ocean), a site with a long record of integrated studies of diffuse vent fluid that document microbial communities composed of Campylobacterota, Aquificota, and methanogenic archaea, carrying out sulfide and hydrogen oxidation, sulfur and nitrate reduction, and methanogenesis (6, 8, 10, 12, 15, 17, 18, 22). In this initial RNA-SIP study, diffuse fluid from one site in one year was incubated with labeled bicarbonate to target autotrophs at mesophilic, thermophilic, and hyperthermophilic growth regimes in the subseafloor (8). Results revealed the power of RNA-SIP in this environment by reducing the complexity of the mixed diffuse fluids, pinpointing the distinct genera and metabolisms that comprised the subseafloor autotrophic community under each growth regime at this site. At 30°C and 55°C, Campylobacterota were dominant, oxidizing hydrogen and primarily reducing nitrate, while methanogenic archaea were the only autotrophs present at 80°C. The method was later used at multiple hydrothermal vent sites along the Mariana back-arc, where RNA-SIP incubations at 55°C and 80°C indicated that autotrophic Campylobacterota were active and present at all but one sampled vent, which instead had a high abundance and activity of autotrophic Aquificota. These differences were attributed to variation in hydrothermal activity due to underlying tectonic setting (23).

Given the ability of RNA-SIP to link specific microbial activities to their metabolisms, genes, and populations across different incubation conditions, we applied this methodology to address hypotheses about the distribution and activity of subseafloor autotrophs in three diffuse flow vents at Axial Seamount: Marker 113, Anemone, and Marker 33. These vents were chosen because they are situated among different vent fields within the Axial caldera, and they are geochemically and microbiologically distinct (10, 27–28). While all of Axial is hosted in basalt, Marker 33 and Marker 113 vent directly from the seafloor lava flows, while Anemone vents through a sulfide mound atop lava flows (10, 27). Fortunato *et al*. (2018) (10) used metagenomics and metatranscriptomics coupled to geochemistry to study the unmanipulated venting fluids from the three sites in three consecutive years. The study found that each site maintained their specific microbial communities and populations over time, but with spatially distinct taxonomy, metabolic potential, and gene transcription profiles. Some organisms and metabolisms were restricted in distribution, such as hydrogenotrophic methanogens, which was linked to host vent chemistry and venting style (10, 28).

Building on this legacy of diffuse fluid research at Axial Seamount, we present two years of RNA-SIP data from these same three diffuse flow vent sites at three temperatures to examine how the active microbial subseafloor autotrophs at Axial Seamount vary with temperature, fluid chemistry, geology of the vent (sulfide vs. basalt), space, and time. We hypothesized that the subseafloor autotrophic microbial population structure at each vent and temperature would be distinct, with transcriptional metabolic profiles reflective of the geological and geochemical characteristics of each individual vent. This study also contributes new *in situ* oxygen data and an expansive metatranscriptomic dataset that enabled differential expression analysis of isotope-enriched transcripts across temperatures within and among vents, and between years. We determined the specific geochemical and thermal conditions under which autotrophic communities and populations are active and identified potential competition amongst these microbial groups for electron acceptors and donors, including oxygen, nitrate, and hydrogen. Overall, these findings expand our knowledge of the distribution and energy acquisition strategies of active subseafloor microbial assemblages in the productive deep-sea hydrothermal vent ecosystem.

## RESULTS

Chemical characteristics of all diffuse flow vents are shown in Tables S1 and S2 and include data from both previously published works (see 10, 28) as well as new oxygen data collected with an *in situ* probe (Figs. 1 and S1, Table S3). At all diffuse flow vents, the temperatures ranged from 18.5 to 28.2°C during sampling, the pH was on average ∼5.6, and the microbial cell concentrations were the same order of magnitude at each site (1.7-6.8 x 10^5^ cells/mL) and all an order of magnitude above concentrations in background seawater (Table S1). However, there were clear differences in the diffuse fluid chemical composition among the three vents (Figs. 1 and S1, Tables S1, S2, and S3). At Marker 113 and 33, NO3^-^ and H2 were depleted, and CH4 concentrations were elevated. At Anemone, there was evidence for depletion of H2. Anemone also had the highest H2S concentrations of all sites. Newly collected oxygen data showed that all the venting sites were depleted relative to near-bottom seawater in the Axial caldera (31 µmol/kg, +/- 2µmol/kg, n=24). Linear regressions of *in situ* O2 against temperature indicate oxygen behaves differently at each site, with zero O2 predicted to occur at 33°C for Marker 113 and at 52°C for Marker 33; at Anemone, regression extrapolates to zero O2 at >100°C (Fig. 1).

**Figure 1.**
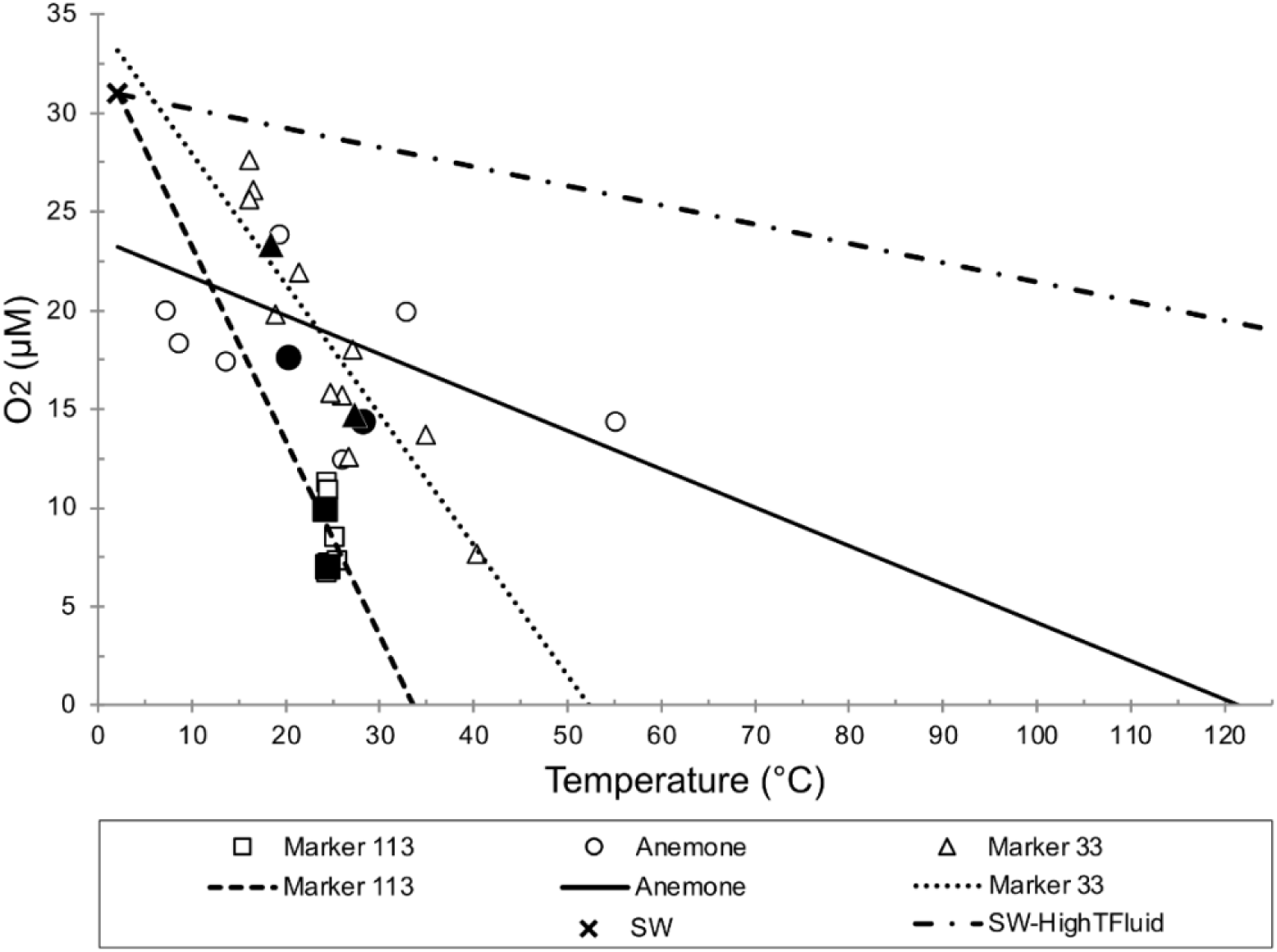
*In situ* oxygen versus temperature for diffuse vents sampled for this study. Near-bottom seawater averages 31 µmol/kg (+/- 2.2) O_2_. Linear regression lines are drawn for each of the three vent sites for the two years sampled, including the ambient seawater point in the regression. The SW-HighTFluid line is between seawater and a hypothetical zero-oxygen end-member at 325°C. Solid data points indicate the exact fluid sampling bag used in the SIP experiment.

### RNA-SIP experiments

In total, 15 sets of RNA-SIP experiments were carried out in 2013 and 2014 using fluids collected from Marker 113, Anemone, and Marker 33 (Tables 1, S4, and S5). Six of these experiments included two timepoints (Table 1). All 15 RNA-SIP experiments showed a heavier buoyant density peak of RNA in the ^13^C-labeled experiment compared to the ^12^C-labeled control, indicating uptake of labeled bicarbonate into RNA (Figs. S3-S5). The highest RNA concentration was chosen from the ^12^C-light and ^13^C-heavy peaks for sequencing. If the experiment had two RNA peaks, the two peak fractions were pooled together and sequenced as one sample (Marker 113 at 30°C and 80°C, Figs. S3C and S5C). For experiments that had two time points, the ^12^C-light and ^13^C-heavy peaks from both time points were sequenced separately. Corresponding ^13^C-light and ^12^C-heavy fractions were also sequenced as additional controls, but given their very low RNA concentrations, the sequencing results were not always successful, and these data were not included in any subsequent analyses.

**Table 1.**
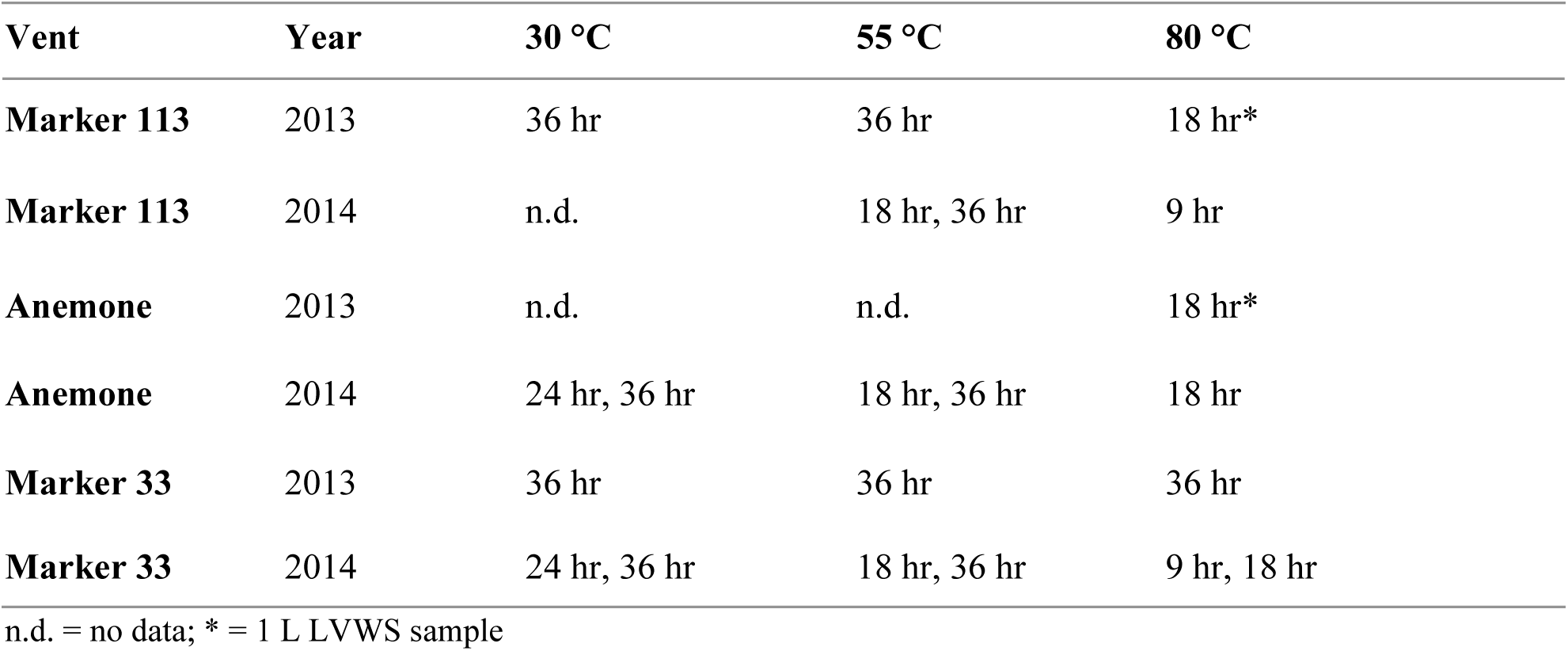
All RNA-SIP experiments, showing vent name, year, temperature of experiment, and time points collected during incubation.

Including technical sequencing replicates, there were 84 total metatranscriptomes (44 ^13^C-heavy and ^12^C-light, 34 ^13^C-light and ^12^C-heavy RNA-SIP experimental metatranscriptomes, and six metatranscriptomes from the *in situ* unmanipulated vent fluid; Table S4) included in this study, totaling over a billion high-quality reads from paired-end Illumina sequencing. Real-time quantitative Reverse Transcription Polymerase Chain Reaction (qRT-PCR) was subsequently carried out to normalize data across experiments using 16S rRNA copies, as shown in Figures S3-S5 (29, 30). In most cases, the highest concentration of RNA corresponded to the highest rRNA copy number.

Results from the ^13^C-heavy RNA-SIP metatranscriptomes are presented as ^13^C-enriched transcripts throughout the manuscript to distinguish these results from the metatranscriptomes of unmanipulated diffuse fluids. As described in the Methods and Supplementary Methods, the ^13^C-enriched fraction of the RNA-SIP metatranscriptomes was annotated and compared with the paired metagenome and metatranscriptome generated from unmanipulated diffuse fluids that were filtered and fixed on the seafloor. This comparison enabled both annotation and interpretation of experimental results and ensured results were representative of the native hydrothermal vent microbial community.

### Taxonomic composition of unmanipulated fluids and ^13^C-enriched fraction of RNA-SIP experiment microbial communities

The unmanipulated fluid metatranscriptomes represent the taxa with detectable transcripts at the instant of sampling via filtration and immediate RNALater fixation on the seafloor. The taxonomy of *in situ* active microbial community members as well as those in the ^13^C-enriched fraction of RNA-SIP experiments was acquired by mapping non-rRNA transcripts to taxonomically classified metagenomic contigs and then normalizing the genera in each sample to 100% (Fig. 2A-D). The taxonomy of the unmanipulated fluids was compared to both the ^12^C-light and ^13^C-enriched RNA-SIP taxonomy, and in all but one experiment, the taxonomies of the ^12^C-light and ^13^C-enriched fractions of the RNA-SIP metatranscriptomes were comparable (Fig. S6). Therefore, results throughout this study will focus only on the ^13^C-enriched fraction metatranscriptomes, as they represent the population of isotope-enriched (active) microbes (Fig. 2).

**Figure 2.**
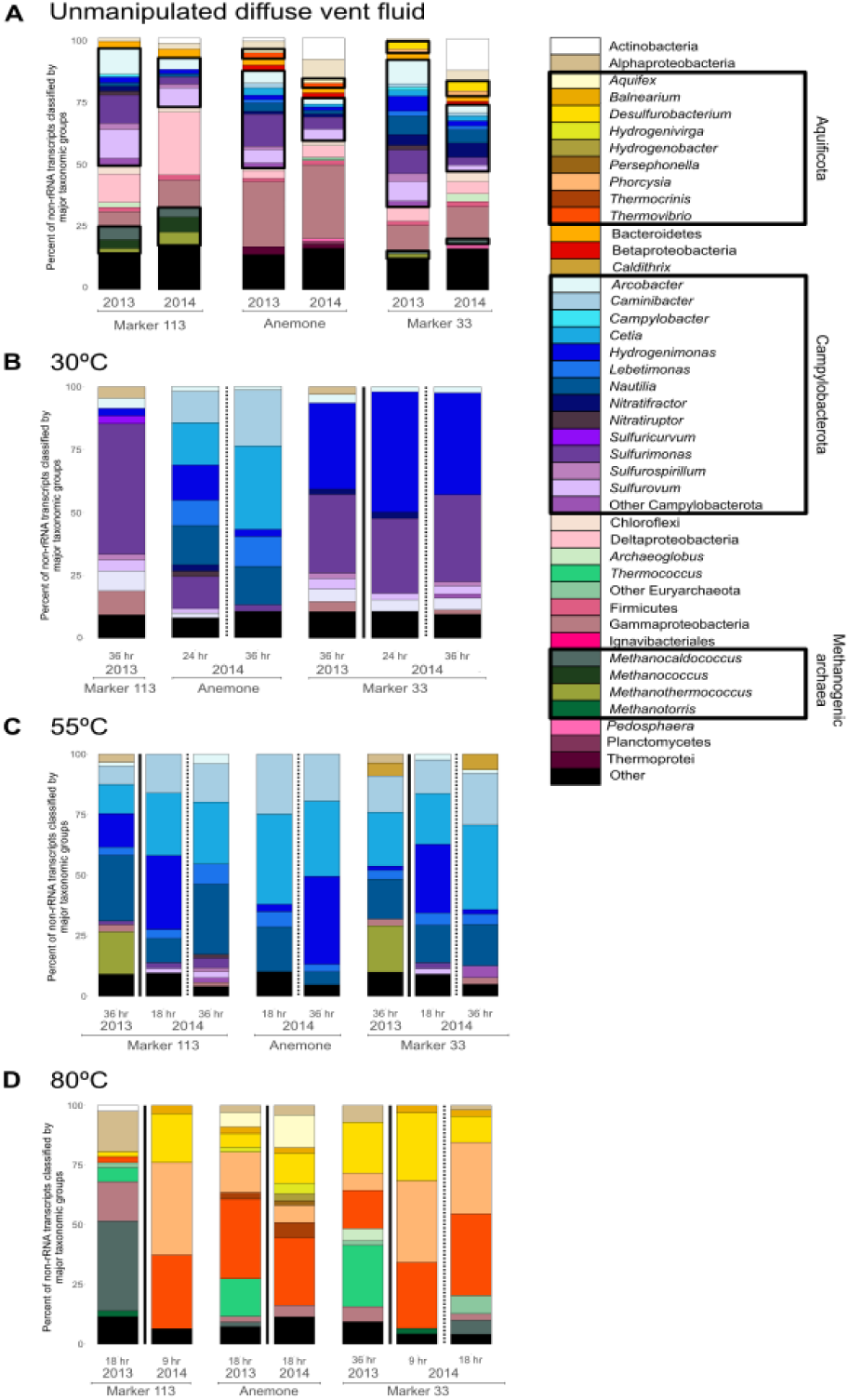
Relative abundance of major taxonomic groups of **A**. Annotated non-rRNA transcripts from unmanipulated fluid metatranscriptomes and annotated non-rRNA transcripts from the ^13^C-enriched fraction of RNA-SIP experiments at **B**. 30°C, **C**. 55°C, and **D**. 80°C. Black boxes indicate the main autotrophic taxonomic groups recovered from the ^13^C-enriched fraction of RNA-SIP metatranscriptomes and are drawn onto 2**A** to show how these taxa make up a small proportion of the unmanipulated fluid metatranscriptome to then become the key autotrophs in the ^13^C-enriched fraction of RNA-SIP metatranscriptomes in 2**B-D.**

All genera active at 30°C in the ^13^C-enriched RNA-SIP metatranscriptomes were also identified in the unmanipulated fluid metatranscriptomes (Fig. 2A). Across all three sites at 30°C, Campylobacterota made up the majority of the active community. *Sulfurimonas* transcripts were observed at all three vents but were most abundant at Marker 113 (52%) (Fig. 2B). Transcripts of *Cetia* (16.8-33.2%) and *Caminibacter* (12.5-22.2%) were found only in the ^13^C-enriched fraction of RNA-SIP experiments from Anemone. At Marker 33, transcripts belonging to *Hydrogenimonas* (34.3%) were more abundant compared to the other vents.

Like the 30°C experiments, all genera active at 55°C in the ^13^C-enriched fraction of the RNA-SIP metatranscriptomes were also identified in the unmanipulated fluid metatranscriptomes (Fig. 2A). Across all three sites at 55°C, the major differences in taxonomy compared to the 30°C ^13^C-enriched fraction of the RNA-SIP experiments, was a shift to higher relative abundances of transcripts from *Cetia, Caminibacter, Nautilia,* and *Hydrogenimonas* (Fig. 2C). These taxa dominated experiments from Marker 113 and Marker 33. However, in 2013, transcripts from thermophilic methanogens were detected in addition to Campylobacterota. *Methanothermococcus* comprised 17.3% and 18.6% of the 2013 ^13^C-enriched fraction of community transcripts at Marker 113 and Marker 33, respectively, but were completely absent at Anemone in 2014.

Unlike at 30°C and 55°C, where Campylobacterota were abundant in both SIP experiments and unmanipulated fluids, many ^13^C-enriched active taxa at 80°C were in low abundance in the unmanipulated fluids (Fig. 2A), including those from the Aquificota and *Methanotorris* at Marker 113, *Phorcysia* at Anemone, and *Thermovibrio* at Marker 33. The major differences in taxonomy in the ^13^C-enriched fraction of 80°C RNA-SIP metatransciptomes across sites included a higher abundance of methanogenic groups at Marker 113 and 33, the presence of *Aquifex* only at Anemone, and persistent detection of *Desulfurobacterium* and *Thermovibrio* at all sites and years (Fig. 2D).

### Metagenome assembled genomes (MAGs)

Binning of the co-assembly of all unmanipulated diffuse fluid, background, and plume metagenomes from Fortunato *et al*. (2018) resulted in 391 bins, with 120 MAGs remaining after filtering for quality (completeness ≥70%, contamination ≤10% based on the presence of single copy bacterial and archaeal marker genes) (10, Table S6). For a phylogenetic tree of the 120 quality-filtered MAGs, see Figure S7. After further filtering these MAGs to remove heterotrophs and those only present in the plume, background, or 2015 metagenomes, 25 MAGs remained. The final 11 MAGs were selected based on having >4% of their coding genome expressed [length of expressed genes(bp)/genome length (bp)] in any individual RNA-SIP ^13^C-enriched fraction (Fig. 3, Tables S6 and S7). The taxonomic composition of the highly transcribed MAGs aligned with ORF-based transcript assignments (Figs. 2 and 3), including groups primarily from the Aquificota, Campylobacterota, and methanogens.

**Figure 3.**
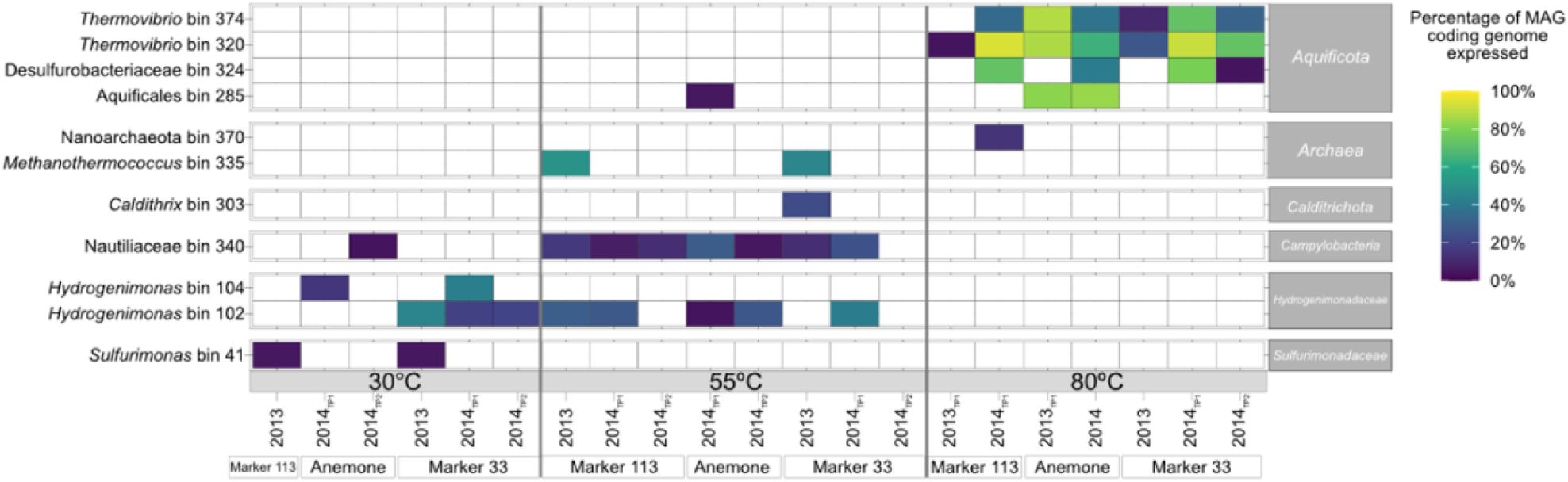
Metagenome Assembled Genomes (MAGs) selected based on having >4% of coding genome expressed in 30, 55 and 80°C ^13^C-enriched fraction of RNA-SIP experiments. Percent of coding genome expressed = length of expressed genes(bp)/genome length (bp).

Both Marker 113 and Marker 33 had *Sulfurimonas* populations (bin 41) transcribed at 30°C (Fig. 3), corresponding to the higher abundance of *Sulfurimonas* transcripts observed at these vents (Fig. 2). Two MAGs classified as *Hydrogenimonas* (bins 102 and 104) were transcribed across different temperatures and vents. Bin 102 was transcribed only at Marker 33 at 30°C, but at 55°C, this bin was transcribed at all vents in at least one timepoint. Bin 104 transcripts, however, were observed only at 30°C at Anemone. Nautiliaceae bin 340 transcripts were ubiquitous at 55°C at all sites and years. *Methanothermococcus* bin 335 transcripts were found at only Marker 113 and Marker 33 at 55°C in 2013, while no methanogen populations were transcribed at Anemone (Figs. 2 and 3).

In general, MAGs expressed a higher proportion of their coding genomes in the 80°C experiments compared to the other temperatures. *Thermovibrio* MAGs (bin 320 and bin 374) were transcribed across all 80°C experiments, matching the high abundance of the genus in the transcripts at all vents (Figs. 2 and 3). The Desulfurobacteriaceae (bin 324) was transcribed in 2014 at all sites at 80°C. Aquificales bin 285, however, was restricted to Anemone only (Fig. 3).

### Functional attributes of ^13^C-enriched active autotrophs

To determine how autotrophs in the ^13^C-enriched fraction of RNA-SIP experiments obtained energy while taking up labeled bicarbonate, we identified transcripts for key genes involved in oxygen, nitrogen, methane, hydrogen, sulfur, and carbon fixation pathways, and compared those to the unmanipulated fluid metatranscriptomes (Fig. 4, Table S8). There were similarities in major metabolic pathways across all sites and samples, but in each year and temperature, the three vent sites differed in magnitude of transcription.

**Figure 4.**
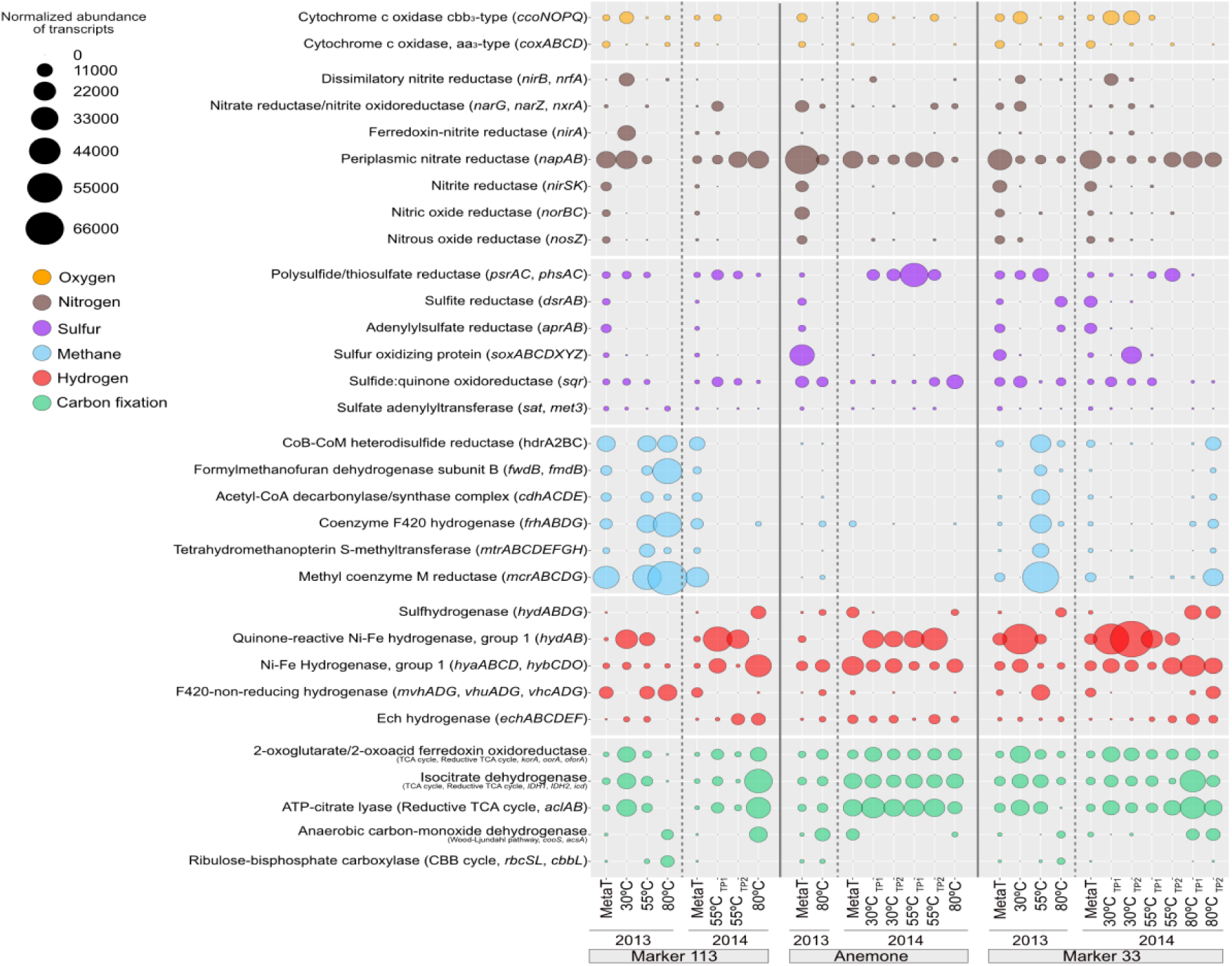
Normalized abundance of non-rRNA transcripts annotated in the unmanipulated diffuse fluid metatranscriptomes (”MetaT“) and ^13^C-enriched fractions of RNA-SIP metatranscriptomes for key marker genes involving oxygen, nitrogen, sulfur, methane, hydrogen, and carbon fixation. Bubble sizes represent the transcripts per million reads (TPM) attributed to each marker gene.

Transcripts for the selected metabolic marker genes were present in most of the unmanipulated fluid metatranscriptomes (Fig. 4), whereas some genes were transcribed in the ^13^C-enriched fraction of the RNA-SIP metatranscriptomes only at certain temperatures and vents, reflecting specific metabolisms being carried out by distinct subseafloor autotrophic populations under different temperature regimes. We also examined the 30 most abundant transcripts at each RNA-SIP experimental temperature (Fig. S8), some of which were metabolic genes that we note here, while other transcripts were related to information processing, replication and repair, and signaling and cellular processes.

Aerobic oxidative phosphorylation gene *aa3*-type cytochrome c oxidase (*coxABCD*) and microaerophily gene *cbb3-*type cytochrome c oxidases (*ccoNOPQ*) transcripts were highest in the ^13^C-enriched fraction of the 30°C RNA-SIP experiments as compared to 55°C and 80°C (Fig. 4). Normalized counts of *ccoNOPQ* peaked at Marker 33 at 30°C in both 2013 and 2014 (7,593–9,608 TPM), attributable to taxa *Aliarcobacter*, *Hydrogenimonas*, *Sulfuricurvum*, *Sulfurimonas*, and *Sulfurovum* (Table S9).

Genes for nitrate reduction and complete denitrification were highly transcribed in almost all the ^13^C-enriched fraction of the RNA-SIP metatranscriptomes (Fig. 4). Of those genes, *napA*, periplasmic nitrate reductase, was the most abundant transcript across all conditions. It was also one of the 30 most transcribed genes at 80°C (1,394–15,195 TPM) (Figs. 4 and S8), predominantly in *Phorcysia* and *Thermovibrio* within the Aquificota. Marker 113, and to a lesser extent, Marker 33, had transcription of genes for dissimilatory nitrate reduction to ammonia (DNRA; *nirAB* and *nrfA*, TPMs 8,131–16,106), and these transcripts were uniquely attributed to *Sulfurospirillum* (Table S9).

Of the six sulfur marker genes selected, *psr* and *phs* are most functional in anaerobic conditions and *sox* in aerobic conditions (31). However, many sulfur metabolism genes can function across diverse oxic conditions, as they catalyze both anaerobic sulfate reduction and the reverse process, sulfur-compound oxidation (32, 33). Anemone had the highest H2S concentrations (Table S1) as well as transcription of genes related to sulfur metabolism compared to the other two vents (Fig. 4). For example, *sqr* had high transcription at 80°C (up to 9,695 TPM from *Aquifex*, *Hydrogenivirga*, and *Phorcysia*), whereas at both Marker 113 and Marker 33, *sqr* was transcribed only at 30°C and 55°C (Fig. 4). The highest transcription of polysulfide/thiosulfate reductase genes *psrAC* and *phsAC* occurred at 55°C in all experiments, with transcripts reaching 27,947 TPM from *Hydrogenimonas*, *Cetia*, and *Nautilia* (Fig. S8). Although *soxABCDXYZ* transcripts were present in unmanipulated fluids at all three vents, only Marker 33’s 2014 ^13^C-enriched RNA-SIP metatranscriptome at 30°C showed pronounced transcription (6,396 TPM), derived from *Hydrogenimonas*, *Sulfurovum*, and *Sulfurimonas* (Table S9).

Methanogenesis genes were transcribed in the unmanipulated fluid samples from Marker 113 and Marker 33 but were largely absent at Anemone (Fig. 4). Transcription of methanogenic metabolic genes was highest in the ^13^C-enriched RNA-SIP metatranscriptomes from Marker 113 in 2013 at 55°C and 80°C (up to 56,765 TPM), and Marker 33 at 55°C in 2013 and at 80°C in 2014 (up to 47,489 TPM). These transcripts were attributed to *Methanothermococcus* and *Methanococcus* at 55°C and *Methanotorris* and *Methanocaldococcus* at 80°C. Methanogenic genes *hdrA2* and *mcrA* at 55°C and *frhA*, *fwdB*, *fmdB*, and *mcrAC* at 80°C were among the 30 most transcribed genes in these temperatures (Fig. S8).

All metatranscriptomes from unmanipulated diffuse vent fluids contained transcripts from selected hydrogenase marker genes (Fig. 4). Notably, throughout all sites and all years, the ^13^C-enriched transcriptional profiles from 80°C RNA-SIP experiments lacked the quinone-reactive Ni-Fe hydrogenase *hydAB*, responsible for transferring electrons from H2 to terminal electron acceptors oxygen, nitrate, sulfate, fumarate, and metals (32). Transcripts for *hydAB* in the 30°C and 55°C experiments belonged to diverse Campylobacterota, predominantly *Sulfuricurvum* (22,883 TPM, Table S9), but also *Sulfurimonas*, *Nautilia*, and *Hydrogenimonas.* Other hydrogen metabolic marker genes, such as sulfhydrogenase (*hydABDG*), energy converting (*echABCDEF*) hydrogenase and Ni-Fe hydrogenase (group 1; *hyaABCD*, *hybCDO*), were in higher abundances at 55°C and 80°C compared to 30°C and associated with *Thermococcus* and Phylum Aquificota (Table S9). F420-non-reducing hydrogenase (*mvhADG*, *vhuADG*, *vhcADG*) used by methanogens was primarily at Marker 113 and 33 at 55°C and 80°C. Some of these hydrogenase marker genes were among the top 30 genes transcribed in the respective temperatures in the dataset (Fig. S8).

Across all ^13^C-enriched RNA-SIP metatranscriptomes at 30°C and 55°C, the most transcribed carbon fixation pathway was the reductive tricarboxylic acid (rTCA) cycle (Fig. 4). The key rTCA marker gene, ATP-citrate lyase *aclAB,* was transcribed in all ^13^C-enriched RNA-SIP metatranscriptomes at 30°C and 55°C, with transcripts attributed to Campylobacterota (Fig. 4, Table S9). At 80°C in both years at Anemone and in 2014 at Marker 113 and Marker 33, *aclAB* transcripts were attributed to Aquificota (Fig. 4, Table S9). In fact, eight genes involved in the rTCA/TCA cycles were among the 30 most abundant genes transcribed at each temperature (Fig. S8). At Marker 113 (2013) and Marker 33 (2014; Tables S8 and S9), in experiments at 80°C when methanogens were active, transcripts for the reductive acetyl-coA (Wood-Ljungdahl) pathway were also abundant, including Wood-Ljungdahl pathway genes (*cooS, acsA*) attributed to *Methanocaldococcus*.

### Differential Expression Analysis of ^13^C-enriched Transcripts

Differential expression analyses using the RNA-SIP ^13^C-enriched transcriptional profiles were conducted to determine statistical differences in transcription across experimental conditions without bias towards any specific functional annotations. In the absence of an untreated control in our dataset—an inherent feature of our manipulation of natural environmental samples—we used a differential expression framework that does not require an untreated control and instead compares expression profiles to an expected expression baseline calibrated to each comparison of interest (34–38). Comparisons were made across temperatures within each vent, across temperatures among the three different vents, and between years within each vent (see Methods and Supplementary Methods; 34-38). Across all ^13^C-enriched RNA-SIP metatranscriptomes, 656 unique genes were significantly differentially expressed (SDE) 1520 times with an adjusted p-value of <0.05 (Table S10). Only genes with a padj < 10^-3^ and log2 fold-change ≥ |2| will be discussed.

Across temperatures within each vent, the highest number of transcripts were SDE (483 unique genes SDE 1030 times, Fig. 5, Table S11). The baseline for this comparison was the average expression of all ^13^C-enriched RNA-SIP metatranscriptomes (n=21; Table 1). Within Marker 113, annotated transcripts with the greatest differences in expression across temperatures included gene transcripts for hydrogen metabolism, nitrate reduction, and carbon fixation (Fig. 5A). At Marker 113, as RNA-SIP experimental temperature increased, the differentially expressed hydrogenase transcripts in the ^13^C-enriched fraction shifted from quinone-reactive Ni-Fe hydrogenase genes overexpressed at 30°C (*hydB*) and 55°C (*hydAB*) and underexpressed at 80°C (*hydAB*), to Ni-Fe hydrogenase (group 1) genes (*hyaB*, *hybC*) overexpressed at 80°C and underexpressed at 55°C. Transcripts for hydrogenase genes *mvhA*, *vhuA,* and *vhcA*, utilized in methanogenesis, were underexpressed at 30°C.

**Figure 5.**
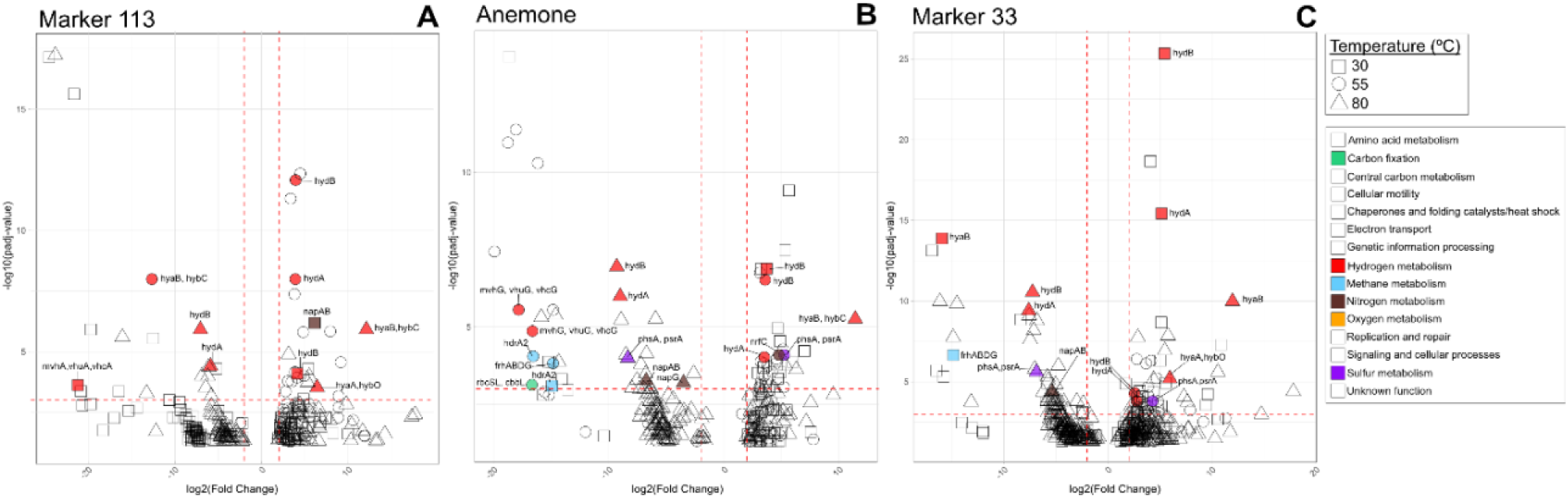
Volcano plots showing the log_2_FC (degree of differential expression) across experimental temperatures at each vent **A.** Marker 113, **B.** Anemone, and **C.** Marker 33. Colored and labeled points correspond to genes involved in carbon fixation, hydrogen, methane, nitrogen, oxygen, and sulfur metabolism that had a padj < 10^-3^ and log_2_FC > |2| [padj; p-value adjusted for multiple tests using the Benjamini-Hochberg procedure to control the false discovery rate (FDR)].

At Anemone, transcribed genes with the greatest differences in expression across temperatures included transcripts related to carbon fixation and hydrogen, methane, nitrogen, and sulfur metabolism (Fig. 5B). At 55°C, transcripts for Calvin-Benson-Bassham (CBB) cycle genes *rbcSL* and *cbbL* were underexpressed. The hydrogen metabolism genes *hydAB*, and *hyaB*, *hybC* followed the same pattern of differential expression as at Marker 113. At all temperatures at Anemone, transcripts for genes involved in oxygen-sensitive methanogenesis (*mvhG*, *vhuG, vhcG; frhABCG*, *hdrA2*) were underexpressed. Nitrate reduction gene transcripts (*napABG*) were underexpressed at 80°C, while transcripts for the nitrite reduction gene *nrfC* were overexpressed at 55°C. Finally, sulfur metabolism gene transcripts for polysulfide/thiosulfate reduction (*phsA*, *psrA*) were underexpressed at 80°C and overexpressed at 55°C (Fig 5B).

There were more differentially expressed transcripts across temperature within Marker 33 than the other vents, reflecting the greater number of metatranscriptomes in the dataset (n=9 ^13^C-enriched RNA-SIP metatranscriptomes, Table 1). These included transcripts for hydrogen, methane, nitrogen, and sulfur metabolism (Fig 5C). Marker 33 had the same pattern for differential expression of hydrogenases *hydAB* at 55°C and *hyaB* at 80°C as both Marker 113 and Anemone.

Transcripts for methanogenesis genes *frhABCG* were underexpressed at 55°C. Marker 33 had the same genes differentially expressed in the same pattern as Anemone for nitrogen and sulfur metabolism.

When comparing across vents by temperature (Fig. S10), 240 transcripts were SDE 329 times, using a baseline expression of normalized transcript counts of all ^13^C-enriched RNA-SIP metatranscriptomes within each temperature (n=6 for 30°C, n=8 for 55°C, and n=9 for 80°C; Table 1). This comparison revealed SDE transcripts for genes related to hydrogen and nitrogen metabolism. At 30°C, nitrate (*napAB*) and nitrite (*nirA*) reduction gene transcripts were overexpressed only at Marker 113, while nitrate/nitrite (*narGZ*, *nxrA*) reduction gene transcripts were overexpressed at Marker 33 (Fig. S10, Table S12). The greatest number of SDE transcripts was observed at 30°C. None were identified at 55°C, and only three transcripts were SDE in the 80°C experiments.

In a comparison between 2013 and 2014 within each vent, 114 unique genes were SDE 161 times (Fig. S11, Table S13). The baseline expression for this analysis was the same that was used for the across vents by temperature comparison, but the contrast in DESeq2 was switched to time instead of temperature (see Methods). Most SDE gene transcripts were overexpressed at Marker 113 and Marker 33 in 2013, including those for a Wood-Ljungahl pathway carbon fixation gene (*cdhACDE*), methanogenesis hydrogenase genes *mvhA*, *vhuA*, and *vhcA* at Marker 33, and several other methanogenesis genes at both Marker 113 and 33. For Figures with SDE genes coded and annotated by additional general functional categories, see Figures S9, S10 (D-F), and S11 (D-F).

## DISCUSSION

In this study, we experimentally manipulated venting fluids to target subseafloor microbes under variable temperature conditions and used stable isotope labeling with inorganic carbon paired with metatranscriptomic sequencing to focus on active autotrophic subseafloor microbes, their metabolic and growth strategies, and their distribution in the subseafloor. While our results broadly corroborated our hypothesis that the subseafloor microbial population structure at each vent and temperature would be distinct, by focusing on only autotrophs at specific temperatures, we also found active, cosmopolitan, subseafloor autotrophs occupying particular thermal niches at all sites, regardless of vent geochemistry or geology. Certain metabolisms and associated lineages such as methanogenic archaea (“specialists”) were restricted in distribution, while other metabolisms geared toward hydrogenotrophic sulfur and nitrate utilization and microaerophily (“generalists”) were widely distributed in many lineages across a wide temperature range, highlighting metabolic flexibility and functional redundancy across diverse taxonomic groups in this ecosystem. Competition for electron donors like hydrogen and electron acceptors like oxygen was apparent in the data, as shifts in taxonomy and transcription highlighted the conditions under which various microbial groups could dominate. The data suggest that while chemical and physical characteristics at each vent control the energy available for microbial growth—as is the case for methanogenesis and sulfide oxidation—some ubiquitous microbial subseafloor lineages within the young oceanic crust at Axial Seamount have adapted to a broad range of subseafloor chemistries and temperatures and are therefore not restricted in distribution.

All ^13^C-enriched fractions of the RNA-SIP experiments experienced a significant distillation of the taxa present in the unmanipulated fluid metatranscriptomes (Fig. 2). Of note is the reduction or disappearance of *Alpha*-, *Delta-*, and *Gammaproteobacteria*, as well as *Actinobacteria*, groups that comprised up to half of the microbial community in unmanipulated fluids and are common in plume and background seawater samples. The RNA-SIP experiments were amended with hydrogen to re-create energy-rich, warm subseafloor conditions, based on both cultivation experiments from Axial (39) as well as failed RNA-SIP experiments without hydrogen enrichment (8). Along with incubation temperature, this likely explains why these common deep ocean taxa were not heavily represented in the ^13^C-enriched RNA-SIP metatranscriptomes, as they were either not adapted to these subseafloor conditions or were heterotrophs. Here, we discuss some of the specific patterns of taxonomic distribution, gene expression, and geochemical features across the three vents to determine how the interplay of fluid chemistry, microbial competition, and functional potential results in the autotrophic activity of microbial populations across vents and temperatures.

### Restricted methanogenesis, widespread hydrogen utilization

Geochemical profiles of diffuse venting fluids consistently indicated biological hydrogen utilization at all sites and between years, but methane production was only evident at Marker 113 and Marker 33 (Fig. S1). These geochemical indicators of biological activity broadly map to the metabolism-focused analyses. ^13^C-enriched transcripts for methanogenesis, an anaerobic metabolism, were restricted to Marker 113 and 33 (Fig. 4). In addition, the top 30 expressed genes (TPM) at 55°C and 80°C included those related to methanogenesis only at these two vents (Fig. S8). This finding also corresponds to the consistent activity of the singular, high quality *Methanothermococcus* MAG (bin 335) at Marker 113 and 33 (Fig. 3). In contrast, at Anemone, methanogenesis-related ^13^C-enriched transcripts were absent in the top 30 expressed genes (Figs. 4 and S8) and were significantly underexpressed (padj < 10^-3^) compared to other sites. This pattern may result from the oxic regime at Anemone that extends into hotter temperatures relative to the other two vents (Figs. 1 and 4).

Whereas methanogenesis was confined to two vents, all sites showed biological utilization of hydrogen, and at Marker 113 and 33, also utilization of nitrate (Fig. S1). ^13^C-enriched transcripts for hydrogen and nitrate utilization were abundant across all conditions, with overexpression of the periplasmic nitrate reductase genes *napAB* as well as genes for quinone-reactive Ni-Fe hydrogenase (*hydAB*), energy-converting hydrogenase (*echABCDEF*), and Ni-Fe hydrogenase (group 1) (*hyaAB*, *hybBO;* Fig. 4) in the differential expression analysis. Taxonomic assignments of transcripts linked hydrogen oxidation coupled with nitrate reduction to diverse genera, including *Sulfurimonas, Cetia, Hydrogenimonas,* and *Caminibacter* at 30°C; *Cetia, Caminibacter, Nautilia, Hydrogenimonas* and those from phylum Aquificota at 55°C; and *Thermovibrio* and *Desulfurobacterium* at 80°C (Fig. 2). Consistent with these transcriptional patterns, diffuse fluid chemistry showed elevated H2S concentrations at Anemone compared to the other two vents (Tables S1 and S2), correlating with high transcription of polysulfide/thiosulfate reduction genes *psrAC* and *phsAC* (up to 14,000 TPM) in tandem with hydrogen oxidation (Fig. 4). Taxa expressing both *psrAC* and *phsAC* along with quinone-reactive Ni-Fe hydrogenase genes (*hydAB*) at Anemone included *Cetia, Nautilia*, and *Hydrogenimonas*; the latter is known to utilize oxygen as a terminal electron acceptor (41).

### Cosmopolitan populations function as subseafloor generalists

Within each year, there were MAGs active in all vents in at least one time point: *Hydrogenimonas* bin 102 (at 55°C in 2014), Nautiliaceae bin 340 (at 55°C in 2013 and 2014)*, Thermovibrio* bin 320 (at 80°C 2013 and 2014), and Desulfurobacteriaceae bin 324 (at 80°C in 2014). The recurrence of active MAGs across vents and temperatures indicated that some vents at Axial Seamount host the same populations of microorganisms, agnostic of vent geology (basalt vs. sulfides) or geochemistry (Fig. 3). Previous studies (e.g. 9-10, 16, 42) determined that established subseafloor microbial communities are mostly restricted by local vent fluid geochemistry and physical structuring of the site. For example, Fortunato *et al*. (2018) showed MAGs from unmanipulated fluid identified as mesophilic and thermophilic methanogens were restricted mainly to Marker 113 and not Marker 33 or Anemone. This result is connected to both vent chemistry (hydrogen, methane concentrations) and physical mixing, allowing for a more stable subseafloor community at Marker 113 compared to other vents (10). However, our results show this may not be true for all lineages at the population level. Our data reveal that some cosmopolitan taxa are capable of colonizing a wide range of environmental niches, functioning as ecosystem generalists, while lineages restricted in distribution, such as methanogens, serve as an example of ecosystem specialists.

The cosmopolitan taxa *Hydrogenimonas* and Nautiliaceae, recovered in 30°C and 55°C experiments, are capable of hydrogenotrophic sulfur and nitrate reduction and are efficient hydrogen oxidizers (43, 44). For example, a study done by Nishimura *et al*. (2010) found that hydrogenases purified from the vent-endemic *Hydrogenimonas thermophila* were phylogenetically related to pathogenic Campylobacterota, known to have high hydrogen oxidation activity (43). Like *Hydrogenimonas, Nautilia profundicola*, another vent-endemic organism, was found to have multiple operons for hydrogenases, including one H2-sensing (Group 2), two H2-uptake (Group 1), and three H2-evolving (Group 4) hydrogenases in their genome. Consistent with the findings from cultured representatives of *Hydrogenimonas* and *Nautilia* that highlight flexibility encoded in their genomes for hydrogen utilization (44), our MAG analyses indicated that bins taxonomically assigned to these genera transcribed three of the four hydrogen marker genes at 55°C, including quinone-reactive Ni-Fe hydrogenase (*hydAB*), Ni-Fe hydrogenase (group 1) (*hyaABCD, hybCDO)*, and energy converting hydrogenase (*echABCDEF*; Table S14), indicating the hydrogenase genes were functionally active in these populations.

Transcripts for hydrogenases were also detected at 80°C at both the community level and in the ubiquitous Aquificota MAGs, supporting the prevalence and importance of hydrogen oxidation in taxa from this phylum. Transcripts for three hydrogenase marker genes, Ni-Fe hydrogenase (group 1) (*hyaABCD*, *hybCDO*), energy-converting hydrogenase (*echABCDEF*), and sulfhydrogenase, were attributed to both *Desulfurobacterium* and *Thermovibrio* (Figs. 2 and 4, Table S9). Desulfurobacteriaceae bin 324 was present at all vents in 2014 and was found to transcribe the hydrogenase maturation gene *hypF* (Table S14). *Thermovibrio* bin 374, found at all vents in 2013 and 2014, transcribed numerous hydrogenase genes, including an F420-non-reducing hydrogenase gene, hydrogenase maturation gene *hypF,* hydrogenase expression/formation genes *hypED*, and multiple sulfhydrogenase subunits (Table S14). The diversity and versatility of these hydrogenase genes may enable these taxa to grow and survive across vents with different chemistries, leading to the cosmopolitan nature of these populations.

### Methanogens and Aquificota compete for hydrogen in subseafloor

Hydrogen is also utilized by hydrogenotrophic methanogens at these vents, and competition for hydrogen among methanogens and the cosmopolitan, hydrogenotrophic sulfur and nitrate reducers of the Aquificota likely occurs in the subseafloor at temperatures >50°C (45–47). Methanogenic growth may be inhibited by growth of these other groups by bringing localized H2 concentrations to a partial pressure below which methanogenesis can occur (48–50). Cultivation experiments showed that *Desulfurobacterium* strain HR11 has a lower H2 requirement and lower *Ks* than *Methanocaldococcus* (30 μM vs. 67 μM; 39, 46, 47). In co-culture experiments with *Methanothermococcus thermolithotrophicus* (Topt 65°C), *Methanocaldococcus jannaschii* (*T*opt 82°C), and *Desulfurobacterium thermolithotrophum* (Topt 72°C), Kubik and Holden (2024) found that *D. thermolithotrophum* outcompeted both methanogens at high and low hydrogen concentrations when grown at a 1:1 ratio of initial inoculum (45). However, when *M. thermolithotrophicus* was grown at a ratio of 10:1 with *D. thermolithotrophum*, the methanogen outcompeted the bacterium, suggesting the conditions under which methanogens can successfully establish first and grow in larger numbers, as observed in the 55°C experiments at Markers 113 and 33, where methanogens were in higher abundance.

An additional factor that can influence the distribution and metabolic strategies of hydrogen-utilizing microbes are physical characteristics of the hydrothermal vents. Building off a reactive transport model developed to examine thermophilic versus hyperthermophilic methanogen growth at Axial vents (28), Kubik and Holden expanded the model with additional hydrogenotrophic autotrophs and determined that when the residence time of subseafloor fluids was long, methanogens dominated, but when fluids had shorter residence times, Aquificota dominated. Lower temperatures combined with longer residence times also favored thermophilic methanogenic growth (45). Taken together, our results support these cultivation and modeling studies and show that in the natural world, methanogens require more specific physicochemical conditions for successful growth, thus limiting their distribution, whereas diverse Aquificota and Campylobacterota are more metabolically flexible and less dependent on geochemistry constrained by fluid residence times, enabling a more cosmopolitan distribution. Anemone’s close (<1 m) proximity to a high temperature (∼300°C) end-member may create short subseafloor fluid residence times; in contrast, Marker 113 and Marker 33 are located farther (∼750 m) from a high-temperature venting source, supporting longer fluid residence times, and thus methanogens (28). Our linked geochemical and meta-omics results suggest that the distribution of some subseafloor microbes may be determined by both a microbe’s efficiency and genetic regulation of hydrogen utilization as well as the interspecies relationships, chemistry, residence time, and temperature of the subseafloor venting fluids.

### Temperature-dependent oxygen regimes structure active autotrophic communities

The novel *in situ* oxygen sensor data enabled new insights into the important role of oxygen as a terminal electron acceptor for subseafloor microbial metabolisms across temperatures and vent locations. Microbial community composition changed with both temperature and site, with Campylobacterota dominating all ^13^C-enriched RNA-SIP metatranscriptomes up to 80°C. At 30°C, ORF-based transcript analysis attributed a substantial proportion of transcripts to *Sulfurimonas* at Marker 113 and *Hydrogenimonas* at Marker 33 (Fig. S12). These results correspond to local oxygen availability: although H2 concentrations were comparable across these two sites, oxygen-temperature trends predicted anoxia at temperatures above 55°C at both Marker 113 and Marker 33 (Fig. 1).

At Marker 113, oxygen concentrations were the lowest of all three vents (7.0 µM in 2013, 9.9 µM in 2014, Fig. 1, Table S1). While fluid residence times and flow rate impact oxygenation, Marker 113’s comparatively low oxygen concentrations may be in part due to *Sulfurimonas* O2 consumption. In the 30°C expression profiles of Marker 113 and Anemone, *Sulfurimonas* transcribed only the cytochrome c oxidase gene, *cbb3-*type (*ccoNOPQ*) and no other genes for alternative electron acceptors, suggestive of a metabolism of oxygen reduction and hydrogen oxidation (Figs. 4 and S12, Table S9). The absence of *Hydrogenimonas* metabolic transcripts and the dominance of *Sulfurimonas* in transcriptional profiles at Marker 113 at 30°C, suggest that microaerophilic *Hydrogenimonas* was limited by low O2 and/or competition with *Sulfurimonas*. By contrast, the metabolic versatility and higher growth optimum of *Hydrogenimonas* likely support its activity at higher temperatures, where it can utilize alternative electron acceptors (51). At Marker 33, *Sulfurimonas* and *Hydrogenimonas* were recovered in equivalent proportions at 30°C. Here, the persistence of oxygen up to 52°C, closer to the growth optimum of *Hydrogenimonas*, may have allowed both taxa to utilize oxygen at higher temperatures at this site (Fig. 1, Table S3). However, these sustained oxygen concentrations may also inhibit colonization by taxa with similar metabolisms, higher optimum growth temperatures, but lower oxygen tolerance, such as *Caminibacter, Cetia, Lebetimonas*, and *Nautilia*.

Beyond the low-temperature patterns, oxygen availability at high temperatures influenced the functional profiles in 80°C incubations at each site. Linear regression modeling showed that oxygen is depleted at 33°C at Marker 113 and 52°C at Marker 33 but persists to at least 100°C at Anemone (Fig. 1). These oxygen models align with the transcript data, as the presence of oxygen at high temperatures at Anemone likely enabled some taxonomic groups to continue utilizing oxygen for microaerophilic sulfide oxidation in 80°C experiments. This result was supported by the transcription of sulfide:quinone oxidoreductase (*sqr*) gene for sulfide oxidation attributed to *Aquifex*, *Hydrogenivirga*, and *Phorcysia*. However, at Marker 113 and Marker 33, the gene for this oxygen-dependent enzyme for autotrophy (40) was not expressed above 55°C, reflecting oxygen depletion at lower temperatures and a shift to alternative electron acceptors such as nitrate.

## Conclusions

This study identified subseafloor microbial primary producers at three spatially separated diffuse vents over two years at Axial Seamount and enabled a granular view of the autotrophic microbes at the base of the deep-sea hydrothermal vent ecosystem. The active autotrophs vary among vents, demonstrating that vent geology, fluid chemistry, and flow path shape the geochemical environments and active metabolisms observed. This dynamic interplay between microbial metabolism and vent fluid chemistry not only influences the community structure but also determines which groups dominate under varying environmental conditions. Hydrogen and oxygen availability and utilization is a key control for the establishment of autotrophs, with competition between autotrophic sulfur-reducing taxa and methanogens for hydrogen evident. These interactions underscore the complex ecological networks at hydrothermal vents, where microbial communities not only adapt to but also actively shape their environment through metabolic exchanges and niche differentiation.

## MATERIALS AND METHODS

### Sample collection

Axial Seamount (Fig. S13) samples were collected using ROVs ROPOS and JASON aboard the *R/V Falkor* and *R/V Thompson* in September 2013 and *R/V Brown* in August 2014 (8, 10). Hydrothermal plume water from 42 m above Anemone (N 45.9335667, W 130.013667) and ambient seawater at 1500 m depth (N 46.27389, W 129.79548) were also collected using a Seabird SBE911 CTD equipped with 10 L Niskin bottles in 2015 aboard the *R/V Thompson*, 5-months post-eruption at Axial Seamount. Water was transferred from Niskin bottles into expandable plastic containers, and 3 L was filtered through 0.22 µm Sterivex filters (Millipore, Billerica, MA, USA) for metagenomic sequencing only.

The Hydrothermal Fluid and Particle Sampler (HFPS; 27) and Large Volume Water Sampler (LVWS; 54) were used to collect fluids at ∼2 cm above the seafloor. The coordinates for each vent are Marker 113: N 45.9227, W 129.9882; Anemone: N 45.9332, W 130.0137; Marker 33: N 45.9332, W 129.9822. Temperature was monitored while drawing fluids using an integrated temperature sensor that was present on both samplers. For sampling with the HFPS and LVWS, a 4 L capacity acid-washed Tedlar bag or a 5 L acid-washed plastic carboy, respectively, was used to collect fluids for RNA-SIP experiments as described in (8). DNA and RNA from diffuse fluids for metagenomic (2013–2015) and metatranscriptomic (2013 and 2014) sequencing were concurrently collected using the HFPS at a rate of 100–150 mL min^-1^ through a 0.22 μm, 47 mm GWSP filter (Millipore) on the seafloor for a total volume of 3 L. All filters were preserved *in situ* with RNALater (Ambion, Grand Island, NY, USA) as described in Fortunato *et al*. (2018) (10). RNA-SIP experiments are reported here only from 2013 and 2014, since there were too few samples from 2015 to include in the analysis, though the unmanipulated diffuse fluid metagenomes were included in the assemblies to provide more phylogenetic and functional coverage representative of the Axial diffuse fluid microbial community.

Hydrothermal fluid samples were collected with the Hydrothermal Fluid and Particle Sampler (HFPS) following procedures described in (27). Vent fluid samples were analyzed on board for pH, total H2S, and dissolved methane and hydrogen. Magnesium and nitrate were analyzed on shore. With the exception of oxygen data, these chemical data were previously reported in (10) and used for methanogen modeling in (28). For this study, HFPS included an oxygen optode (SeaBird SBE 63) capable of measuring oxygen concentration in fluids up to a maximum of 40°C. Temperature was monitored at the sample intake, near the sensor block, and inside the oxygen sensor housing. Oxygen measurements were made just before or after collecting fluid samples, with the intake in the same position to replicate collected water samples. Water was pulled from the titanium/Teflon sampling manifold through the sensor, with some cooling taking place along the flow path. The SBE63 optode does not require a constant flow rate, does not consume oxygen, and can make measurements when flow is stopped. The SBE63 sends 1 Hz serial data, which is received by the HFPS on-board computer. Oxygen concentration is calculated using the temperature measured in the sensor housing, the ambient pressure and salinity (derived from ROV sensors and input manually to the software), and the factory calibration, which covers a temperature range from 2°C to 30°C. All the measurements for this study had temperatures at the sensor within the calibration range. Over the 2013-2015 period of this study, the average oxygen concentration of near-bottom seawater within the caldera, where all diffuse vents for this study are located, was 31 µmol/kg (+/- 2µmol/kg, n=24, Fig 1, Table S2). Oxygen increases significantly with increasing depth at Axial Seamount, where the summit is below the oxygen minimum.

### RNA-SIP experimental procedures

Carboys and Tedlar bags were recovered from the ROV and brought to the shipboard lab, where filling into evacuated bottles began immediately. Both were kept at ambient temperature during filling and remained sealed to minimize oxygen exposure, although some likely occurred. The Tedlar bag was housed in a water filled plastic cylinder for collection. As the bag was emptied during bottle filling, additional water was pumped into the cylinder to keep pressure on the bag and minimize outgassing. The time between ROV recovery and experimental set-up was less than 1 hour. For LVWS samples in 2013, fluid from carboys was pumped using a peristaltic pump into evacuated 1 L Pyrex bottles with air-tight butyl stoppers to a volume of 1060 mL. For all other samples in 2013 and 2014, fluid was pumped using a peristaltic pump from the sampling bags or carboys into evacuated 500 mL Pyrex bottles with air-tight butyl stoppers to a volume of 530 mL.

Due to limitations in the amount of fluid able to be collected (4 to 5 L), no biological replicates were conducted. During filling, steps were taken to minimize introduction of oxygen and outgassing of fluid. Prior to filling, ^13^C-labeled sodium bicarbonate (Cambridge Isotope Laboratories, Tewksbury, MA, USA) for the experimental condition or ^12^C-labeled sodium bicarbonate (Sigma, St. Louis, MO, USA) for the control condition was added to each bottle to a final concentration of 10 mM over background. After filling, the pH of each bottle was measured using pH paper and then adjusted to vent fluid pH (<6.5) by adding 1–2.5 mL of 10% HCl. The headspace was filled with pure H2 gas using a syringe in the quantity of 20 mL (1698 μmol/kg) for each 500 mL bottle and 60 mL (2547 μmol/kg) for the 1L bottles. ^12^C- and ^13^C-bicarbonate amended bottles were each incubated at 30, 55, and 80°C for various times as shown in Table 1 in the main text. Incubation times were chosen to ensure incorporation of the labeled bicarbonate and to minimize cross-feeding, as initially determined in Fortunato and Huber (2016). However, after observation of cross-feeding in these initial experiments, timepoints were added and incubation times reduced for further RNA-SIP experiments. After incubation, the fluid in each bottle was filtered through 0.22 μm Sterivex filters (Millipore). Filtration was completed in less than 30 min, and Sterivex filters were immediately preserved in RNALater (Ambion) and frozen at −80°C.

RNA from SIP experiments was extracted according to (8) and detailed in the Supplementary Methods. Following RNA extraction and quantification, each RNA-SIP experiment required isopycnic centrifugation to separate the heavy RNA (labeled with ^13^C—the active autotrophs) from the light RNA (members of the community that did not take up the ^13^C label, representing the less active members of that community) as described in (8) and the Supplementary Methods.

Four metatranscriptomic libraries were constructed for each RNA-SIP experiment: two from the ^12^C-control (^12^C-light and ^12^C-heavy) and two from the ^13^C-experiment, (^13^C-light and ^13^C-heavy) as described in (8) except for four samples: the second timepoints in 2014 of Marker 113 at 55°C, Marker 33 at 30°C and 80°C, and the only timepoint at Anemone at 80°C. The ^12^C-control incubation peak(s) of highest RNA concentration were used to construct the ^12^C-light library, which represents the natural community. Similarly, the ^13^C-experiment incubation peak(s) of highest RNA concentration were used to construct the ^13^C-heavy library, which represents the labeled, active autotrophic community. The other two libraries from each temperature, ^12^C-heavy and ^13^C-light, were constructed to act as additional controls but were not taken through full bioinformatic processing. In some cases (denoted in Figs. S3-S5), two RNA fractions were combined for library construction.

### Metagenomic bioinformatic analyses

All metagenomic libraries were constructed as described in the Supplementary Methods. There were 11 metagenome samples processed, including metagenomes from all three vent sites in 2015 (Table S4; 10). These samples were included to provide a comprehensive functional repertoire in diffuse fluid communities at Axial Seamount (10). A pipeline with code used in metagenomic processing can be found here: https://github.com/corporeal-snow-albatross-5/Axial_Metagenomics. Samples were first trimmed of Illumina adapter sequences using Trimmomatic (v0.32; 55), quality checked using FastQC (v0.11.9; 56), and quality results were aggregated across samples into a single report using MultiQC (v1.12; 57). These initial trimming and quality control steps were done using the qc-trim Snakemake pipeline (https://github.com/shu251/qc-trim). The separate forward reads and separate reverse reads were combined using the cat (concatenate) command in Unix and then concatenated forward and reverse read files were combined using the fq2fa (v1.1.3) function within the IDBA_UD assembler (58; https://github.com/loneknightpy/idba). Each of the metagenomes were assembled using IDBA_UD (v1.1.3), using a max kmer size of 120 (58; https://github.com/loneknightpy/idba). Contig and scaffold quality was evaluated using MetaQUAST (v5.0.2; 59), and the best assembly given steps of 20 from kmer lengths of 20-120 was chosen for further analysis. Each metagenome was submitted to the DOE Joint Genome Institute’s Integrated Microbial Genome Expert Review (IMG/ER) for annotation. IMG/ER JGI GOLD Analysis Project ID numbers can be found in Table S4. Open reading frame (ORF) information from the IMG/ER annotation was extracted using a python script to create .ffn files from .gff (containing contig and locus identity, start and stop sites of ORFs, and strand information) and .fna files (containing the nucleotide sequence of each ORF). All metagenome .ffn files were then concatenated to make one long, annotated assembly using the cat command in Unix.

The ORFs predicted in each metatranscriptomic (mRNA) file were mapped back to the concatenated metagenomic assembly, which was first indexed using the kallisto index function within the pseudoaligner Kallisto prior to mapping (v.0.46.0; 60). Post-mapping, the KEGG Ontology (KO) ID’s and the taxonomy of each predicted ORF in the metagenomes were retrieved from IMG/ER. Cutoffs on the minimum requirements of an e-score of 10^-10^, 30% amino acid identity, and alignment length of 40 amino acids were imposed, as used in (10). Additionally, eukaryote and virus hits were filtered out of the taxonomy file. Before filtering, there were 5,432,185 contigs classified as eukaryotes or viruses, and after filtering, the classified contig count was 5,309,746. Therefore only 2.25% of the total contigs were identified as eukaryotes or viruses, meaning most hits belonged to bacteria and archaea. The filtered KO tables of each metagenome were concatenated with each other, and the same was done for the taxonomy tables. The concatenated KO and taxonomy tables were then combined with each other using an inner_join in the R dplyr package (61, 62), based on target_id (predicted ORF).

### Metagenome Assembled Genome (MAG) generation and analysis

In addition to an annotated, concatenated assembly, a co-assembly using the quality-filtered and trimmed reads from 11 metagenomic samples was also constructed using MEGAHIT (v1.2.9; 63) with parameters --min-contig-len = 200, --presets = “meta-sensitive”. Assembly statistics were checked using MetaQUAST (v5.0.2; 59). The resulting assembly had 2,318,625 contigs that were at least 500 bp in length; of those contigs, the N50 = 721 bp and maximum length = 280,659 bp. The co-assembled metagenome (hereafter referred to as the “co-assembly”) was annotated through the DOE Joint Genome Institute’s Integrated Microbial Genome Expert Review (IMG/ER) pipeline. IMG/ER JGI GOLD Analysis Project ID numbers can be found in Table S4. The resulting gene calls, .gff annotation file, and phylodist taxonomy of contigs, were used as the reference material for the metagenome assembled genomes (MAGs), differential gene expression analyses, taxonomy assessments, and overviews of functional potential versus expression. KEGG Ontology (KO) ID’s and the taxonomy of each predicted ORF in the co-assembly were retrieved from IMG/ER. Cutoffs on the minimum requirements of an e-score of 10^-10^, 30% amino acid identity, and alignment length of 40 amino acids were imposed, as used in (10).

MetaBAT2 (v2.15; 64) was used to generate binned contigs. The co-assembly was first indexed using the bwa index within BWA mem (v7.17; 65) and then metagenomes were mapped to the indexed file. The resulting output was processed through Samtools (v1.13; 66). The .sam file generated from mapping was converted to a .bam file using the samtools view function and resulting .bam files were sorted using samtools sort. Prior to binning, depth (also called coverage) files were created for each metagenome and binned with a minimum contig length of 1500 using MetaBAT2. DAS Tool was then used to resolve redundancy among bins and provide an optimized bin set (v1.1.7, 67). Bins were taxonomically classified using GTDB-tk (v2.1.1; 68).

We refer to unscreened binning outputs as ‘bins.’ Bins with >70% completeness and <10% contamination are termed metagenome-assembled genomes (MAGs; Table S5). Genomic binning from all the diffuse fluid metagenomes resulted in 391 bins, and after removing redundant bins using DAS Tool (v1.1.7; 67), 120 MAGs remained with 45 having <10% contamination and being >70% complete (Table S5). The MAGs were further filtered to remove known heterotrophs (based on literature searches) and MAGs solely sourced from the plume, background, or 2015 metagenomes. This resulted in 25 MAGs remaining. To further narrow the MAGs to those most active in our samples, we determined the percent expressed coding genome for each bin within each of the RNA-SIP metatranscriptomic samples. The expressed coding genome was calculated as the length of the transcripts recruited to each bin (bp) for each sample, divided by MAG size (bp). The final 11 MAGs were selected based on having >4% of their coding genome expressed in any individual RNA-SIP ^13^C-enriched metatranscriptome (Fig. 3 in the main text, Table S7).

### Metatranscriptomic bioinformatic analyses

Metatranscriptomic libraries from unmanipulated fluids were constructed as described in the Supplementary Methods. In total, there were 84 metatranscriptome samples processed: 6 metatranscriptomes from unmanipulated fluids at each vent in 2013 and 2014 and 78 RNA-SIP metatranscriptomes (Table S4). A pipeline containing the code used to process metatranscriptomes is located here: https://github.com/corporeal-snow-albatross-5/Axial_Metatranscriptome. Metatranscriptomic sequences were trimmed and quality checked with the same methods and software parameters as the metagenomic sequences. The separate forward reads and separate reverse reads were concatenated using the cat command in Unix and then concatenated forward and reverse read files were combined using Fast Length Adjustment of Short reads (FLASH; v.1.2.11; 69), which is a program used to combine fastq files. The only parameter set was for read length, which was 110 bp. Although metatranscriptome libraries were constructed with selective amplification for mRNA (not rRNA), the SIP metatranscriptomes were not. To separate the rRNA for preliminary taxonomic analyses from the mRNA for downstream processing, SortMeRNA (v4.2.0; 70) was used (Table S5). The SILVA v.132 bacterial and archaeal 16S (id85), 23S (id98), and 5S (id98) databases were used as reference databases to map the rRNA. The rest of the parameters were: --fastx for a fastq output format, --other for mRNA reads that did not map to the databases, --out2, and --pairedout for non-aligned reads. The percentage of reads mapping to rRNA from SortMeRNA can be found in Table S5.

Fastq reads from SortMeRNA rRNA output were converted to fasta format using the fq2fa (v.1.1.3) function within IDBA_UD assembler (58; https://github.com/loneknightpy/idba). rRNA taxonomy was classified using mothur classify.seqs function and the SILVA (v.132) 16S rRNA reference database (71). The 5S rRNA database used as one of the reference databases for rRNA mapping in SortMeRNA also contains eukaryotes. As such, the eukaryote hits were removed before further processing. Taxonomy count tables were generated using R (61), and bar plotting for rRNA was done via ggplot2 (72) within tidyverse (73) in R. Resulting plots were only used to compare to ORF transcript taxonomy and are not included in this paper.

Fastq mRNA reads were mapped to the indexed concatenated metagenomic assembly using the kallisto quant function within Kallisto (v0.46.0; 60), which also normalized all ORFs that mapped, providing Transcripts Per Million (TPM) values for each ORF. TPM normalizes for both gene length and depth of sequencing, allowing for accurate comparisons between samples (74). Sample names were attached to kallisto output files, then all files were concatenated and tpm values of 0 were removed. This table was then attached to the KO and taxonomy table made from the IMG/ER output using an inner_join in the R dplyr package (61, 62), based on target_id (predicted ORF). 784,486 total ORFs from all samples joined out of 4,027,381 total, which means that 80.521% of ORFs from the concatenated metagenome were unmatched in KO and taxonomy in the metatranscriptomes and SIP metatranscriptomes. There were no eukaryote hits in the ORFs that had a match with both KO and taxonomy. There were only 11 virus hits in these samples of 784,486 total ORFs that did have matching KO’s and taxonomy. Because few viruses and no eukaryotes were found, they were removed for further processing and to enable comparability between the rRNA and ORF taxonomy. All figures and statistical analyses were completed in R (61).

### Differential expression analysis

Metatranscriptomic (mRNA) samples from the ^13^C-enriched fractions of Marker 113, Anemone, and Marker 33 incubations at 30, 55, and 80°C from 2013 and 2014 were pseudoaligned to the indexed co-assembly using kallisto (parameters: --single -l 151 -s 41) (v0.46.0; 60). The kallisto outputs of metatranscriptome pseudocounts were tabulated with the R package tximport (v1.26.1; 36) using a custom annotation database of unique genes collapsed into protein-coding gene representatives. Due to the diversity in metagenomes, the co-assembly annotation contained multiple targets for gene products. To reduce complexity in functional expression analyses, protein-coding genes observed in the co-assembly were indexed into protein families using IMG’s highest confidence annotation such as KEGG, COG, pfam, and TIGRFAM. ORFs annotated with only “hypothetical protein” were discarded. The annotation database indexed 2,428,812 genes representing 14,942 unique genes collapsed into protein-coding gene representatives observed in the co-assembly. When referring to genes and proteins in our evaluations of differential functional expression, we are using this consolidated database.

Differential expression analyses were conducted in R with DESeq2 (v1.38.3; 34, 35), and code can be found here: https://github.com/corporeal-snow-albatross-5/Axial_Differential_Expression. Our experimental design included three different conditions (vent, temperature, and year) with two or more levels (the possible values each variable can take in the DE statistical model). For vent, there are three levels: Marker 113, Anemone, and Marker 33; for temperature there are also three levels: 30°C, 55°C, and 80°C; for year, there are two levels: 2013 and 2014. Because the number of samples across conditions was uneven (i.e. the number of RNA-SIP experiments differed between years and across vents and temperatures, Table 2), we implemented methodology for imbalanced designs and multi-group data, as described in Love *et al*. (2014b) (35). First, we manually defined contrasts (grouping of samples for each comparison) to account for an imbalance of samples in the DESeq2 model design (37, 38). This allowed for the model to identify differential gene expression without requiring an untreated control. Subsetting the dataset minimized the observed within-group variability caused by unparameterized interaction of variables, thereby reducing stochasticity and improving assessment of significant genes. This permitted us to tailor baseline gene expression for each of the three comparisons: across experimental temperatures within each vent (1030 significantly differentially expressed (SDE) transcripts; Table S11), across vents by temperature (329 SDE transcripts; Table S12), and between 2013 and 2014 within each vent (161 SDE transcripts; Table S13).

For temperature comparisons within each vent (i.e. all Anemone experiments at 30, 55, and 80°C compared to one another), the baseline expression for genes was the normalized expression of all ^13^C-enriched metatranscriptomic fractions (n=21; Table 2). For across vent comparisons at a single temperature (i.e. All 30°C experiments across all vents), as well as comparisons between years within each vent (i.e. 2013 Marker 113 80°C incubation compared to 2014 Marker 113 80°C incubation), gene baseline expressions were calculated at each incubation temperature: 6 ^13^C-enriched metatranscriptomes at 30°C; 8 ^13^C-enriched metatranscriptomes at 55°C; and 7 ^13^C-enriched metatranscriptomes at 80°C. Volcano plots show genes with p-values adjusted for multiple test correction using the Benjamini-Hochberg procedure to control the false discovery rate (FDR, 35), with padj < 0.001 and log2 fold-change > |2|; only genes involved in carbon fixation, hydrogen, methane, nitrogen, oxygen, and sulfur metabolism are shown in Figures 6, S10A-C, and S11A-C. Plots with expanded gene function annotations are shown in Figures S9, S10D-F, and S11D-F. For additional details about how the differential expression analysis was conducted, see the Supplementary Methods.

## DATA AVAILABILITY

All raw metagenomic and metatranscriptomic read data are publicly available through the Sequence Read Archive (SRA). Metagenomic assemblies are publicly available through JGI GOLD. All accession numbers are located in Table S4.

## Supporting information

Main Figures

## ACKNOWLEDGEMENTS

This work was funded by the Gordon and Betty Moore Foundation Grant GBMF3297; the NSF Center for Dark Energy Biosphere Investigations (C-DEBI) (OCE-0939564), contribution number TBD, NOAA/PMEL, contribution number TBD; the Joint Institute for the Study of the Atmosphere and Ocean (JISAO) under NOAA Cooperative Agreement NA15OAR4320063, contribution number TBD; and the National Aeronautics and Space Administration (NASA) grant number 80NSSC19K1427.

The NOAA/PMEL supported this work with ship time in 2014 and through funding to the Earth Ocean Interactions group. The 2013 data collected in this study includes work supported by the Schmidt Ocean Institute during cruise FK010–2013 aboard R/V Falkor. The NOAA/PMEL (through funding to the Earth Ocean Interactions group) and GBMF supported field work in 2014 with ship and vehicle time aboard the R/V Brown (RB2014-03). We thank the captains and crews of the *R/V Falkor, R/V Thompson* and *R/V Brown* as well as the outstanding ROV teams of the vehicles *ROPOS*, *SuBastian*, and *JASON*.

Sarah Hu, Connor Scannerton, and Margrethe Serres provided guidance on the analysis and code used to process this dataset, while the Keck Facility at the Josephine Bay Paul Center at the Marine Biological Laboratory provided support for sequencing. Kevin Roe, Noah Lawrence-Slavas, Susan Merle, Andra Bobbit, and William Chadwick provided support at sea.

