## Supplementary Figures for "Metabolic and Population Profiles of Active Subseafloor Autotrophs in Young Oceanic Crust at Deep-Sea Hydrothermal Vents"

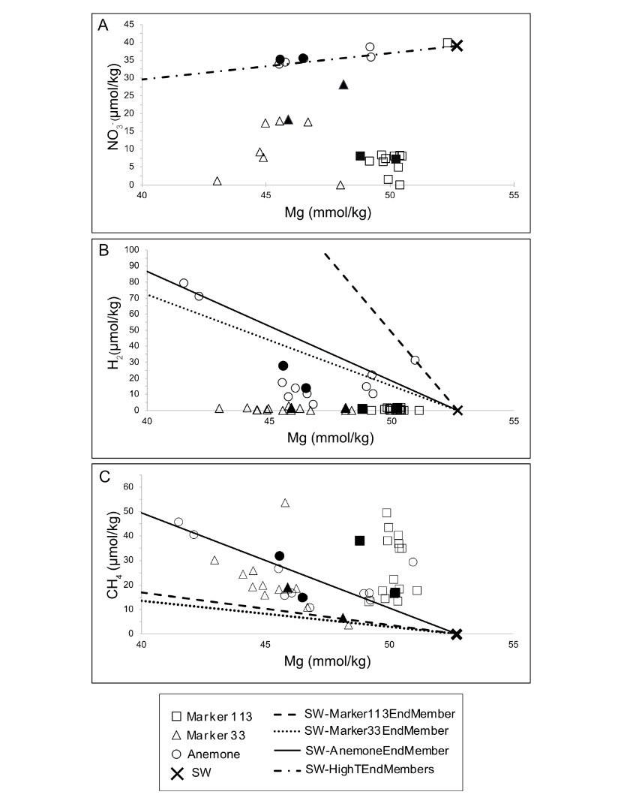

**Figure S1.** Vent chemical species measured at diffuse vent sites Marker 113, Anemone, and Marker 33 in 2013 and 2014. Regression lines are drawn from seawater to end-member fluid concentration for each vent and chemical species to represent conservative mixing between seawater and the high-temperature fluid for **A.** NO_3_^-^. All vent end-members had 0 NO_3_^-^, so the single regression line is between a 0 Mg/0 NO_3_^-^ end-member and seawater. **B.** H_2_ and **C.** CH_4_ concentrations. Solid data points indicate the exact fluid sampling bag used in the SIP experiment.

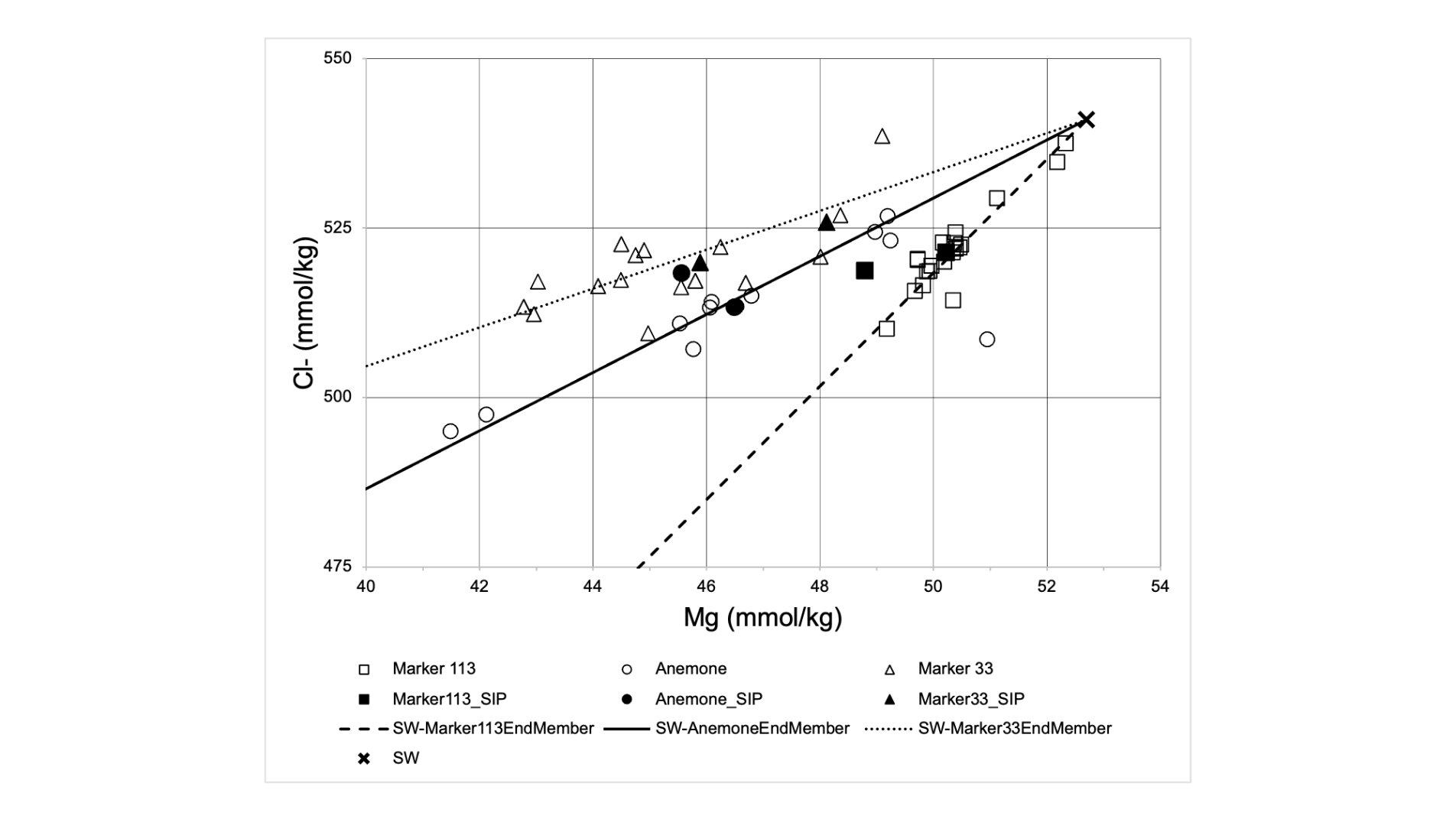

**Figure S2.** Chloride vs. magnesium for diffuse vents in this study. Data points are all samples taken from 2013 through 2015. End-member chloride concentrations are 100 mmol/kg for Marker 113, 315 mmol/kg for Anemone, 390 mmol/kg for Marker 33. Chloride concentrations were used to extrapolate endmember H_2_ and CH_4_ concentrations, as described and reported in Stewart et al., 2019 (28).

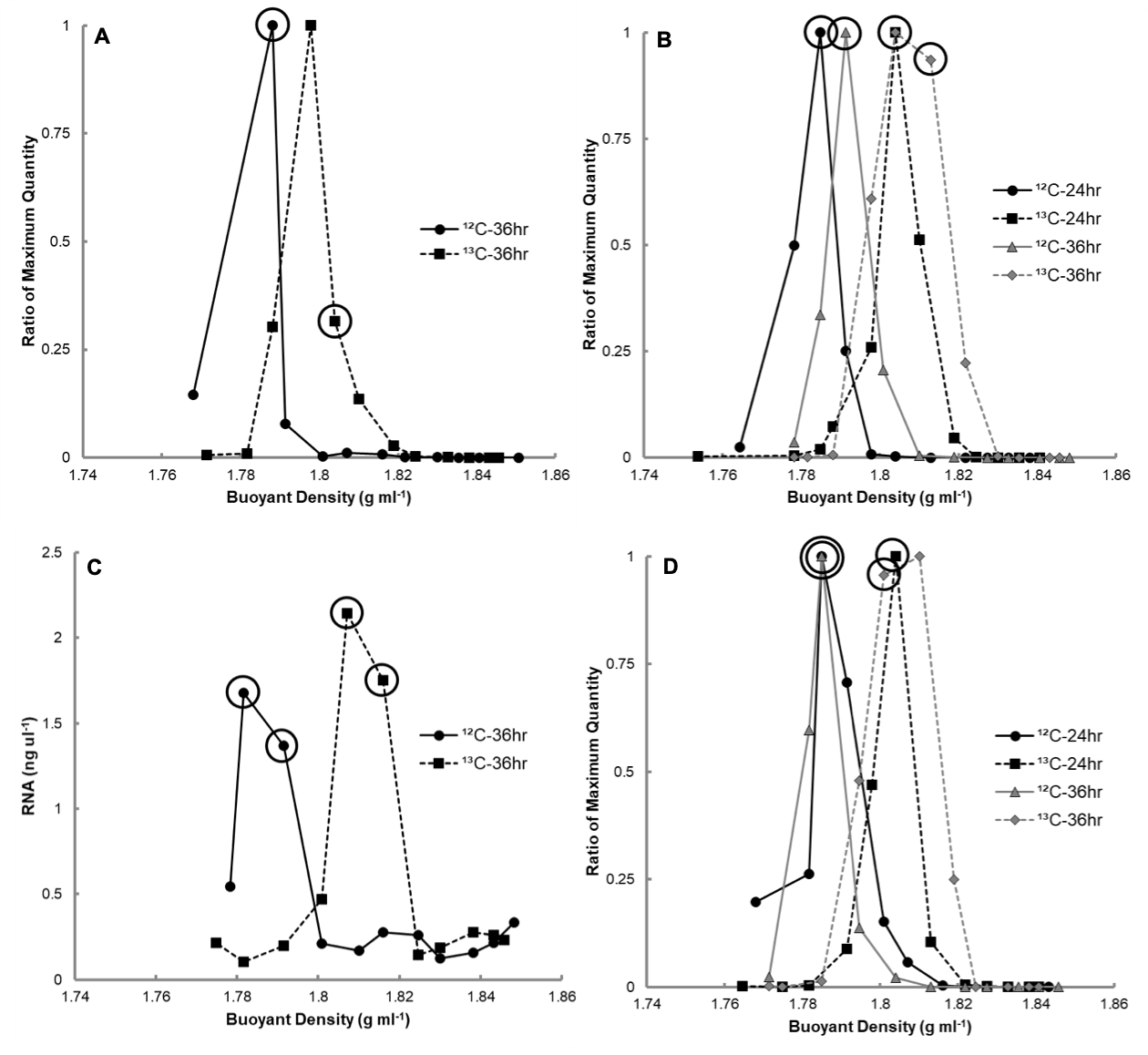

**Figure S3.** rRNA abundance in density gradient fractions of 30°C RNA-SIP experiments for **A**. Marker 33 (2013); **B**. Marker 33 (2014); **C**. Marker 113 (2013); and **D**. Anemone (2014). The buoyant density (grams per milliliter) of each fraction is depicted on the x-axis, and the amount of 16S rRNA determined by RT-qPCR is on the y-axis. The amount of 16S rRNA is displayed as the ratio of the maximum quantity in order to compare results between RNA-SIP experiments. For panel **C**, total RNA concentration is shown. Circles indicate fractions sequenced for metatranscriptomes.

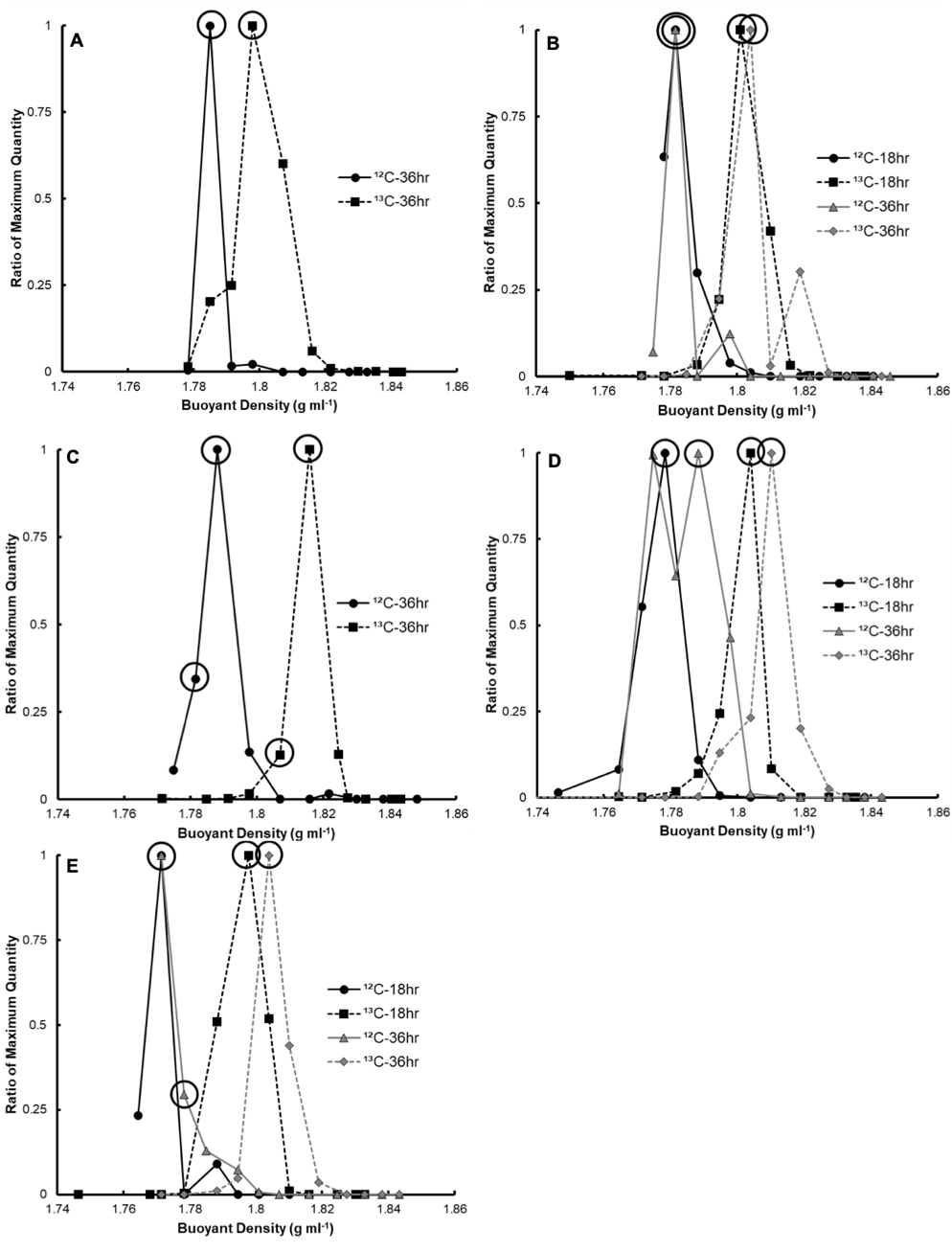

**Figure S4.** 16S rRNA abundance in density gradient fractions of 55°C RNA-SIP experiments for **A.** Marker 33 (2013); **B.** Marker 33 (2014); **C**. Marker 113 (2013); **D.** Marker 113 (2014); and **E.** Anemone (2014). The buoyant density (grams per milliliter) of each fraction is depicted on the x-axis, and the amount of 16S rRNA determined by RT-qPCR is on the y-axis. The amount of 16S rRNA is displayed as the ratio of the maximum quantity in order to compare results between RNA-SIP experiments. Circles indicate fractions sequenced for metatranscriptomes.

 
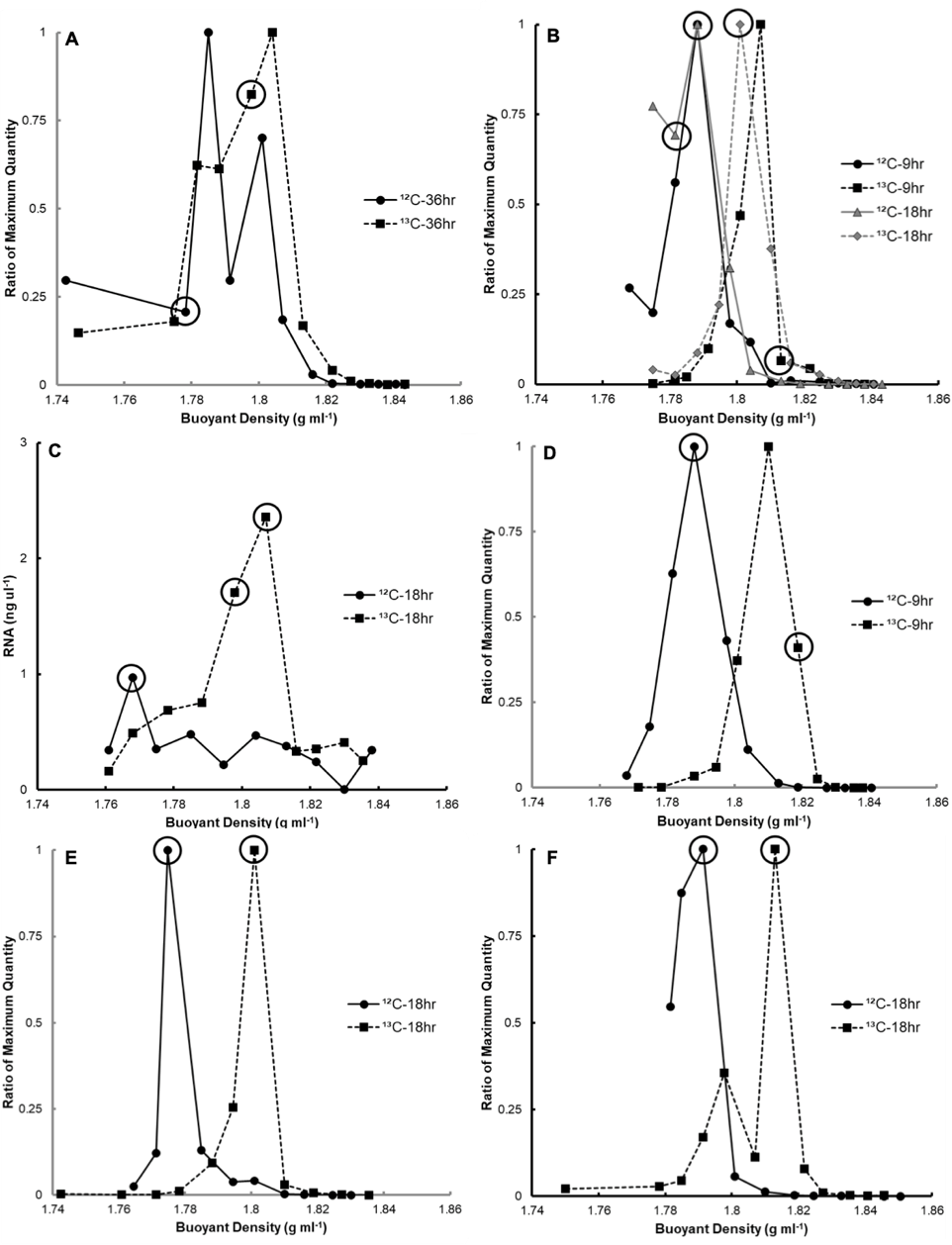

**Figure S5.** 16S rRNA abundance in density gradient fractions of 80 °C RNA-SIP experiments for **A**. Marker 33 (2013); **B.** Marker 33 (2014); **C.** Marker 113 (2013); **D.** Marker 113 (2014); **E.** Anemone (2013); and **F.** Anemone (2014). The buoyant density (grams per milliliter) of each fraction is depicted on the x-axis, and the amount of 16S rRNA determined by RT-qPCR is on the y-axis. The amount of 16S rRNA is displayed as the ratio of the maximum quantity in order to compare results between RNA-SIP experiments. For panel **C**, total RNA concentration is shown. Circles indicate fractions sequenced for metatranscriptomes.

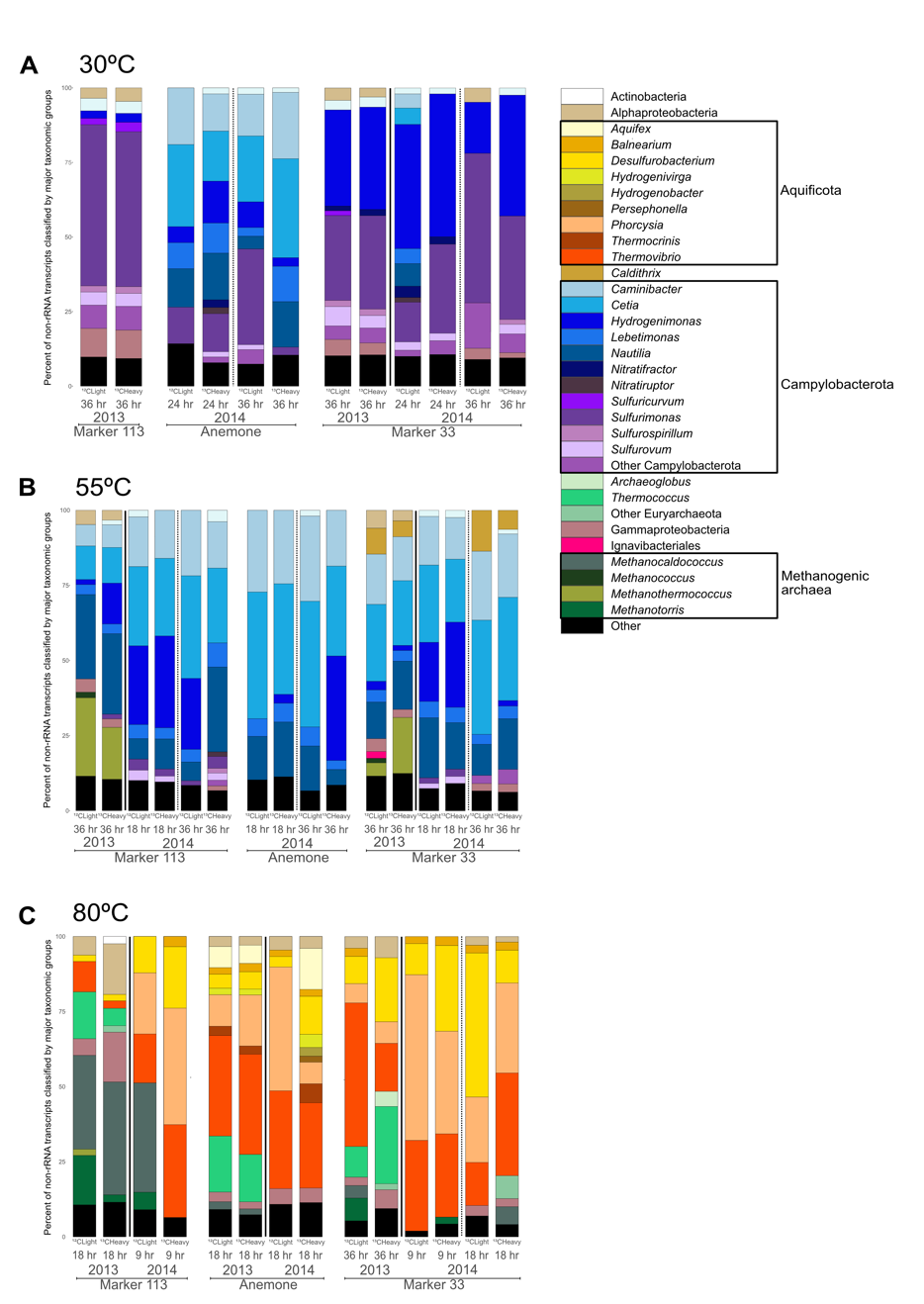

**Figure S6.** Relative abundance of major taxonomic groups of annotated non-rRNA transcripts from the ^12^C-light control fraction and ^13^C-enriched fraction of RNA-SIP experiments at **A**. 30°C, **B**. 55°C, and **C**. 80°C. Black boxes in the key are drawn around the Aquificota, Campylobacterota, and methanogenic archaea indicate the main autotrophic taxonomic groups recovered from the RNA-SIP metatranscriptomes.

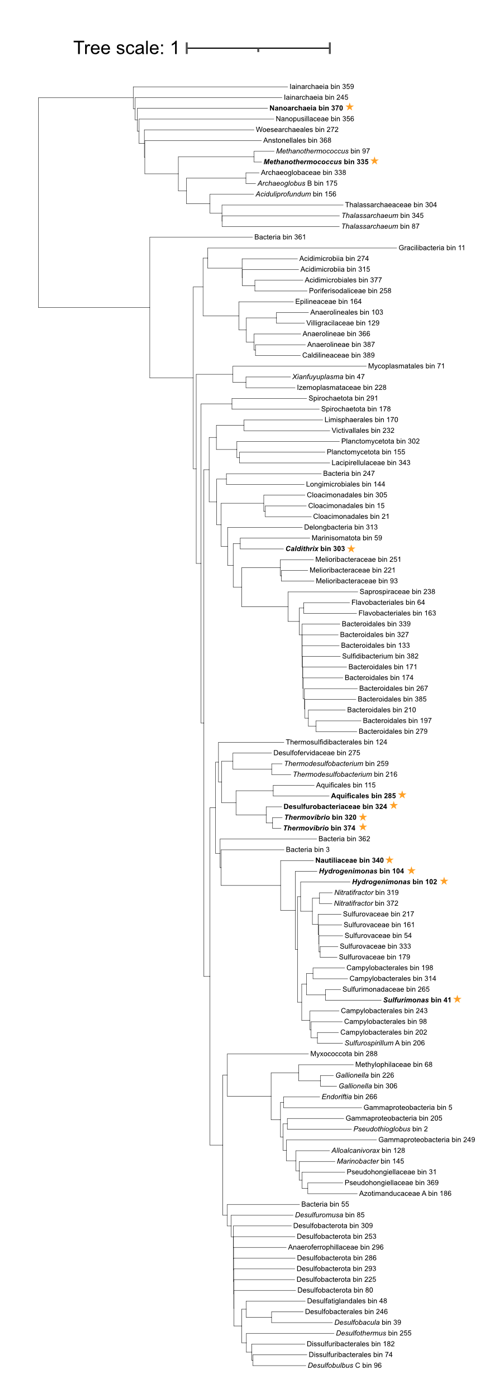

**Figure S7**. Neighbor-joining phylogenetic tree of the 120 quality-filtered metagenome-assembled genomes (MAGs) recovered with >70% completion and <10% contamination. Stars denote the 11 MAGs discussed in the main manuscript with >4% of their coding genome expressed.

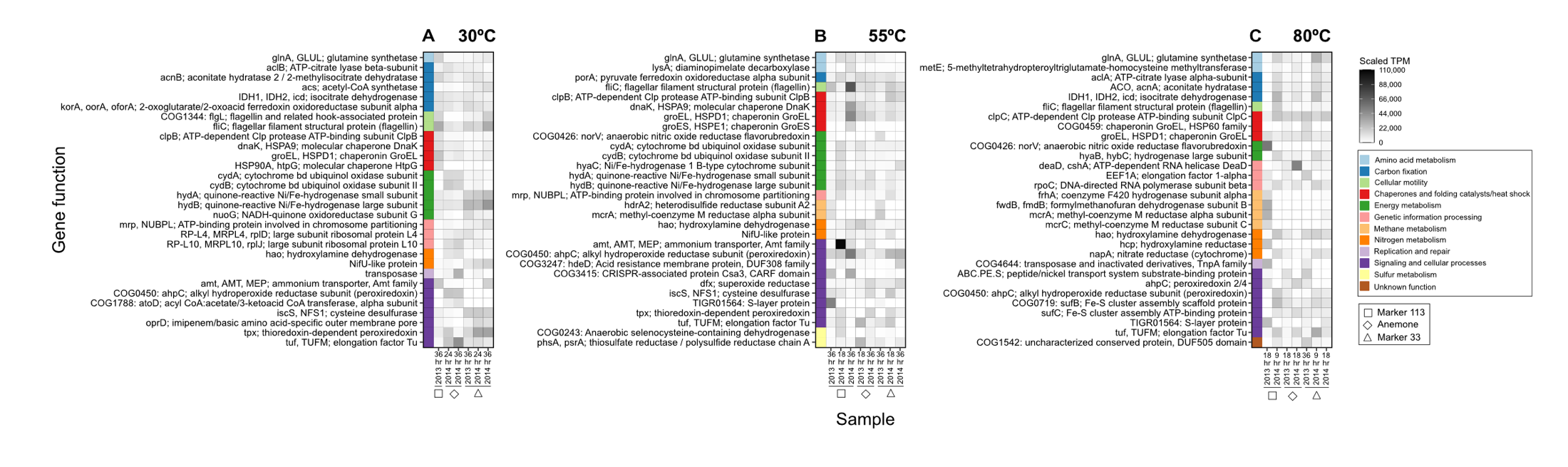

**Figure S8**. Top 30 expressed genes by transcripts per million (TPM) across the ^13^C-enriched RNA-SIP metatranscriptomes at **A**. 30°C, **B**. 55°C, and **C**. 80°C.

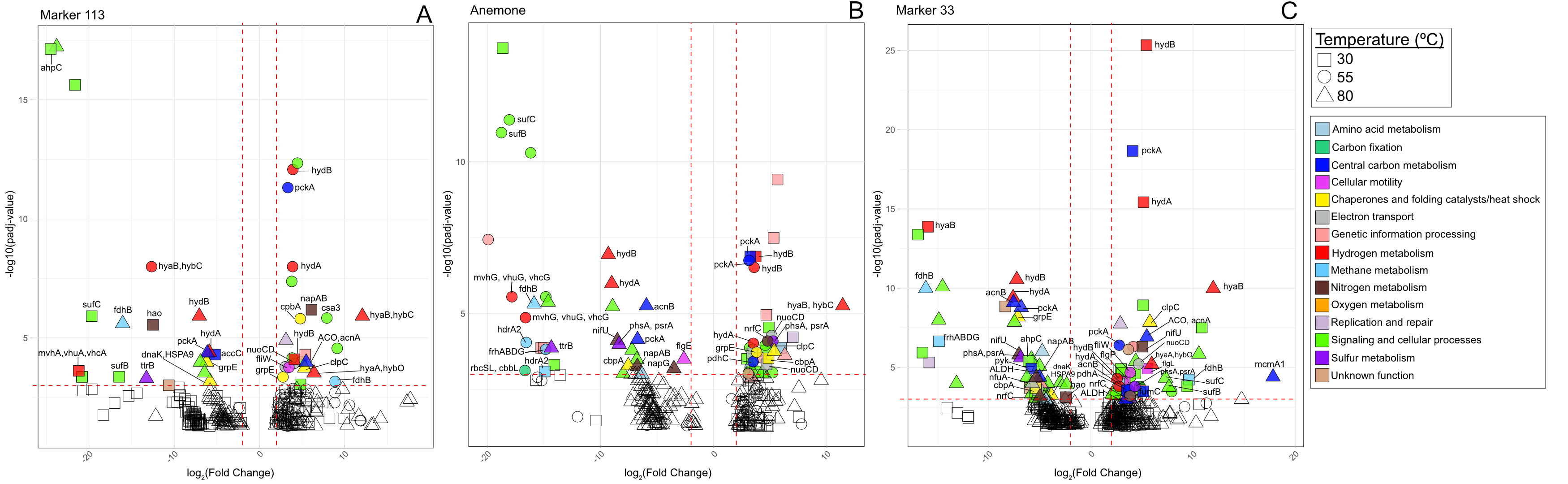

**Figure S9.** Volcano plots showing the log_2_FC (degree of differential expression) across experimental temperatures within each vent at **A.** Marker 113, **B.** Anemone, and **C.** Marker 33. Colored points correspond to all genes that had a padj < 10^-3^ and log_2_FC > |2| (padj; p-value adjusted for multiple tests using the Benjamini-Hochberg procedure to control the false discovery rate (FDR)). Only genes involved in carbon fixation, central carbon metabolism, cellular motility, heat shock, electron transport, Fe-S cluster assembly, CRISPR/Cas system proteins and hydrogen, methane, nitrogen, oxygen, and sulfur metabolism are labeled with gene names.

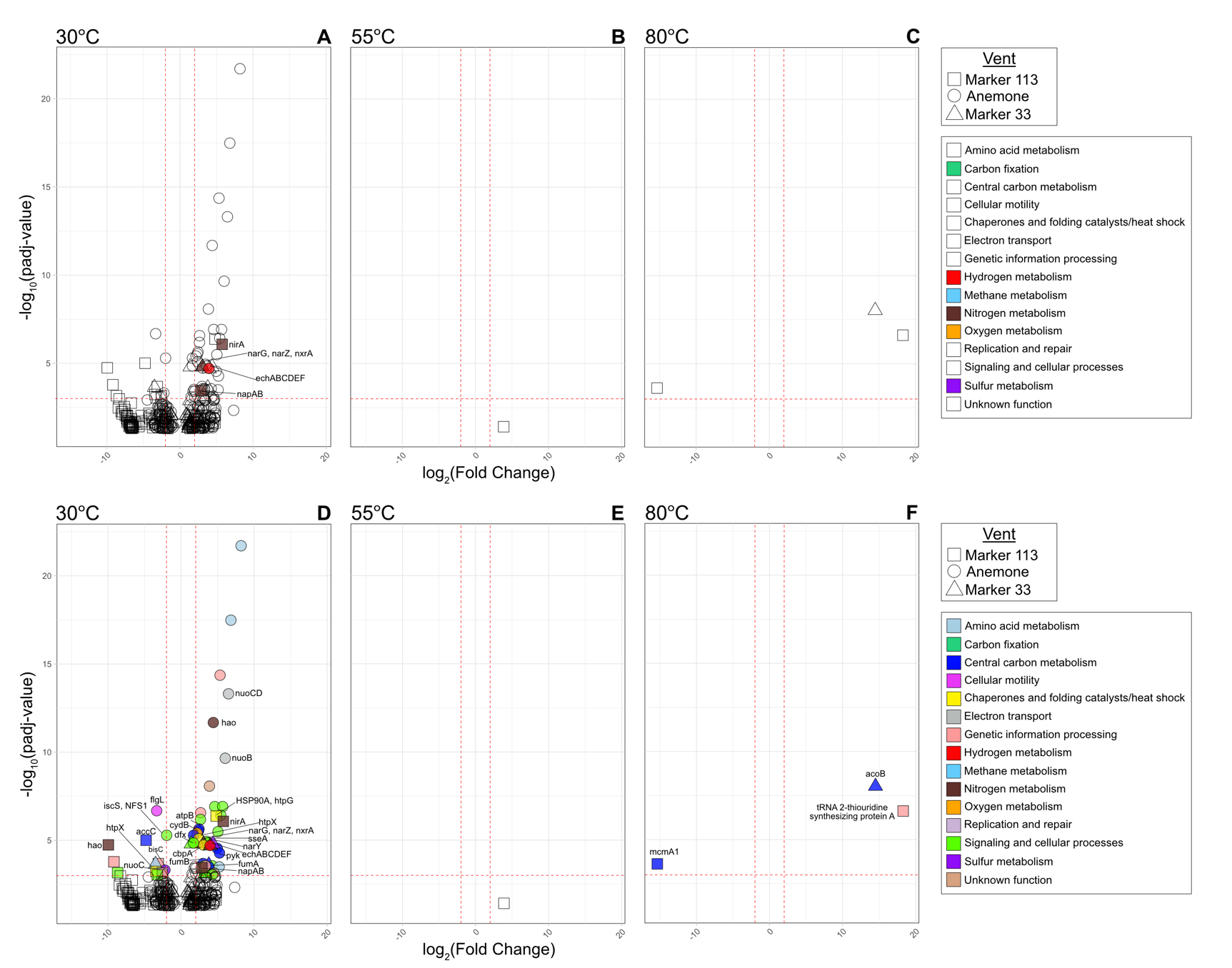

**Figure S10.** Volcano plots showing the log_2_FC (degree of differential expression) across vents by experimental temperature at **A, D.** 30°C, **B, E**. 55°C, and **C, F**. 80°C. Top panel (**A**-**C**), highlights genes involved in carbon fixation and hydrogen, methane, nitrogen, oxygen, and sulfur metabolism with log₂FC > |2| and padj < 10⁻³ (Benjamini-Hochberg adjusted). Bottom panel (**D**-**F**) additionally labels genes related to central carbon metabolism, motility, heat shock, electron transport, Fe–S cluster assembly, and CRISPR/Cas systems.

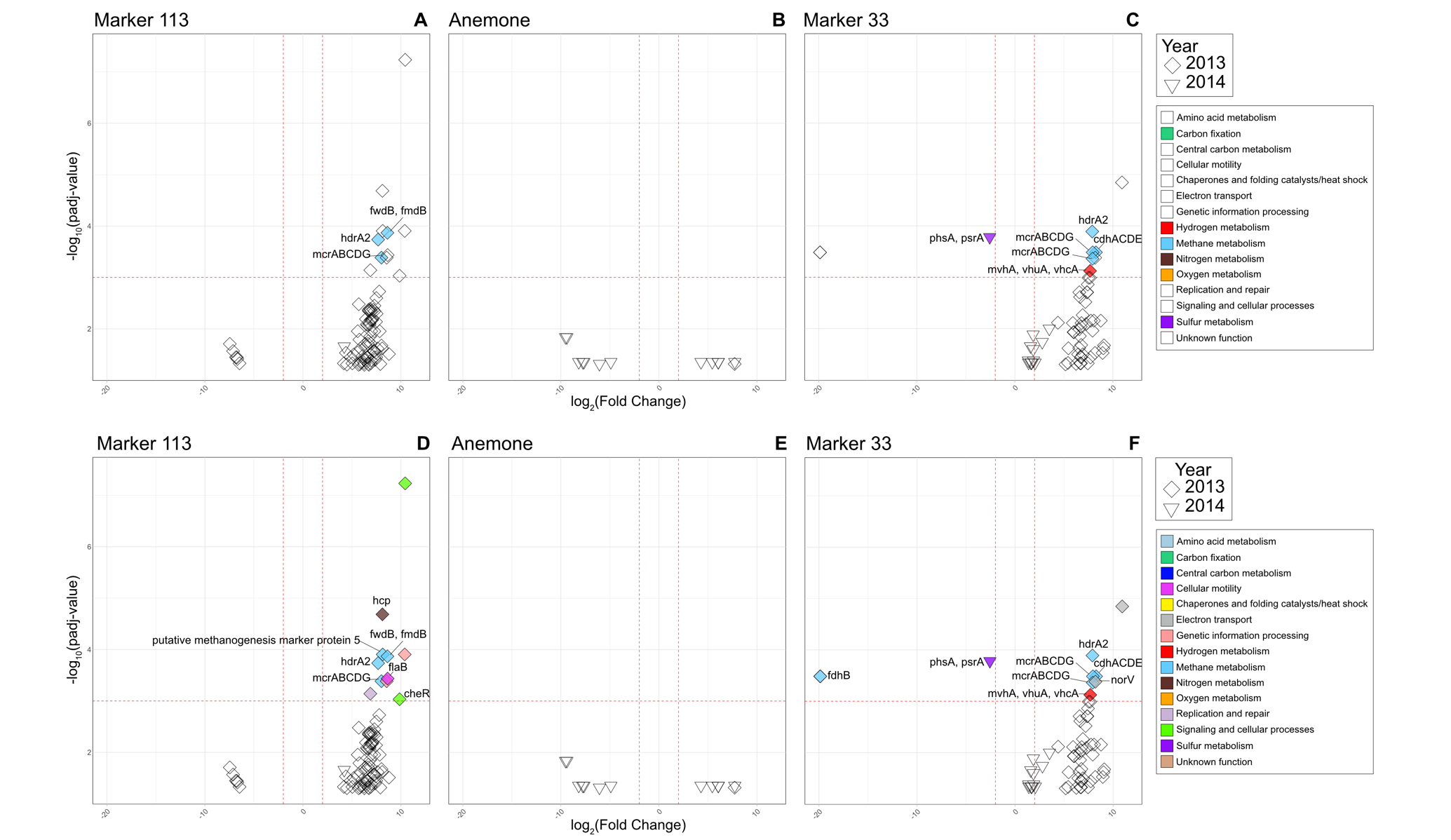
**Figure S11.** Volcano plots showing the log_2_FC (degree of differential expression) between 2013 and 2014 within each vent at **A, D.** Marker 113, **B, E**. Anemone, and **C, F**. Marker 33. Top panel (**A**-**C**), highlights genes involved in carbon fixation and hydrogen, methane, nitrogen, oxygen, and sulfur metabolism with log₂FC > |2| and padj < 10⁻³ (Benjamini-Hochberg adjusted). Bottom panel (**D**-**F**) additionally labels genes related to central carbon metabolism, motility, heat shock, electron transport, Fe–S cluster assembly, and CRISPR/Cas systems.

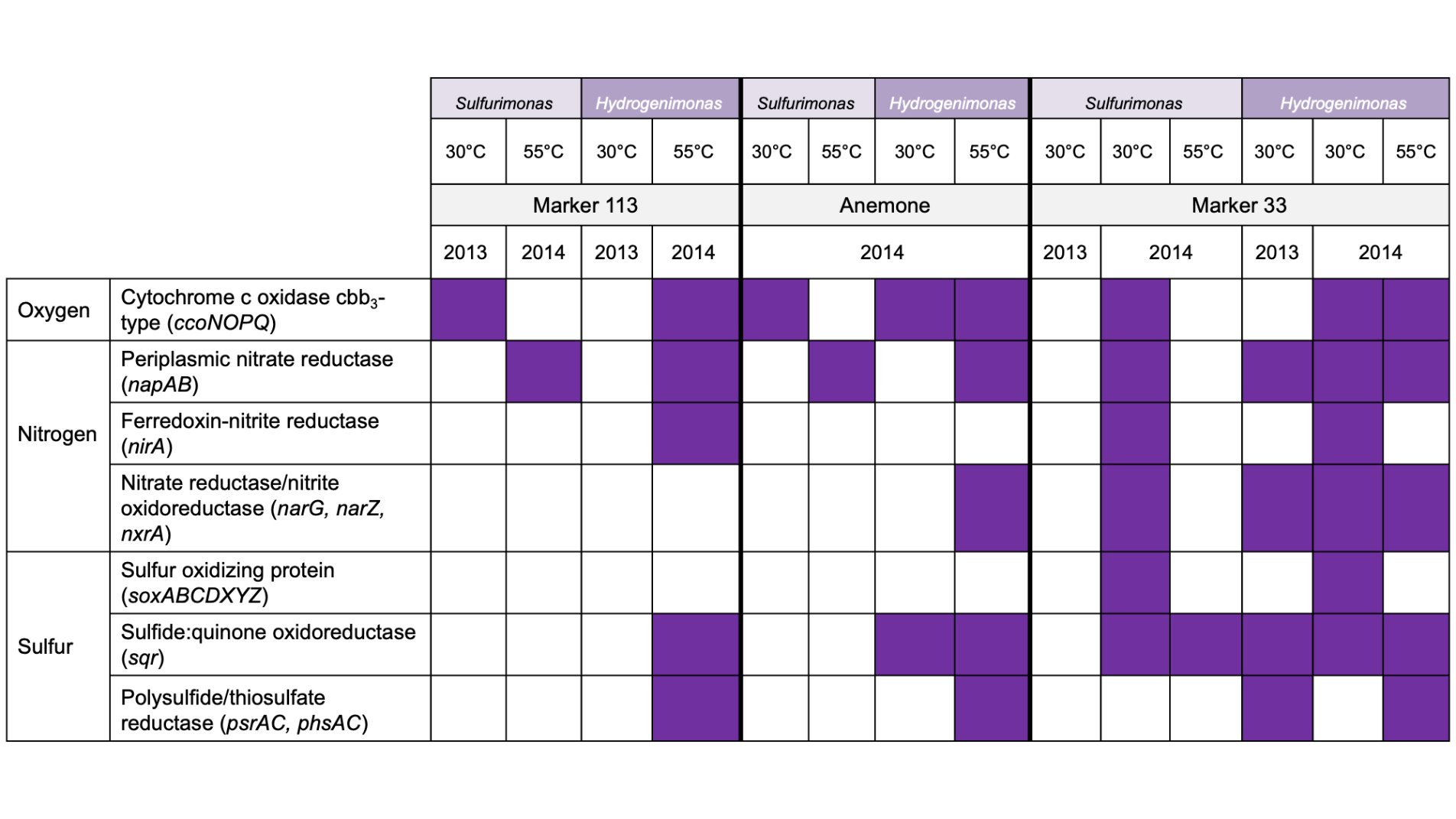

**Figure S12.** Presence/absence plot of dissimilatory genes for oxygen, nitrogen, and sulfur metabolism transcription by *Sulfurimonas* and *Hydrogenimonas* in the ^13^C-enriched RNA-SIP metatranscriptomes at 30 and 55°C.

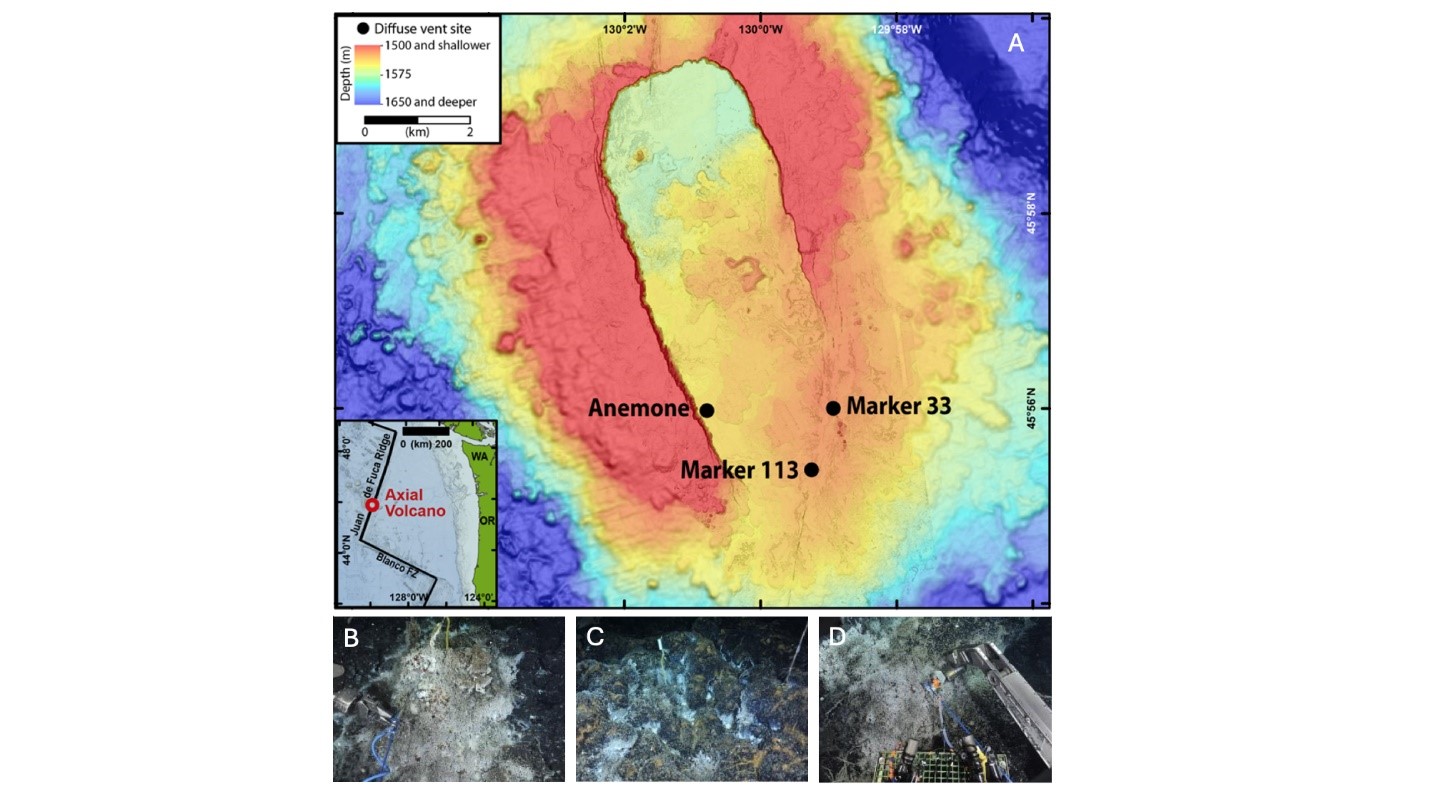

**Figure S13.** **A.** Map of Axial Seamount showing locations of the three diffuse vents sampled, **B.** Marker 113, **C.** Anemone, and **D.** Marker 33. Map from Fortunato *et al*., 2018 (10). Images from ROV Jason (WHOI).
