## Supplementary Methods for "Metabolic and Population Profiles of Active Subseafloor Autotrophs in Young Oceanic Crust at Deep-Sea Hydrothermal Vents"

**RNA Extraction from Sterivex filter**

RNA from SIP experiments was extracted according to (8) by first cutting the Sterivex filter into pieces with a sterile razor blade then following the protocol in the mirVana miRNA isolation kit (Ambion). Biological material was removed from the filter pieces with a bead-beating step using RNA PowerSoil beads (MoBio, Carlsbad, CA, USA), followed by an extraction with acid phenol:chloroform. A total volume of 100 μL RNA was extracted and any remaining DNA was removed using the Turbo-DNase kit (Ambion).

**Isopycnic centrifugation, fractionation and qRT-PCR**

Following RNA extraction and quantification, each RNA-SIP experiment required isopycnic centrifugation to separate the heavy RNA (labeled with ^13^C—the active autotrophs) from the light RNA (members of the community that did not take up the ^13^C label, representing the less active members of that community) as described in (8). To prepare the gradient solution for each sample, 5.1 mL of CsTFA (~2 g mL-1; GE Healthcare Life Sciences, Piscataway, NJ, USA), 185 μL formamide, and a mixture of 500-750 ng RNA and gradient buffer solution (0.1 M Tris-HCl, 0.1 M KCl, 0.1 mM EDTA) up to 1 mL were first mixed in a 15 mL falcon tube prior to centrifugation (25). Once mixed, the refractive index was measured for each sample to ensure a median RI of 1.3729 +/- 0.0002, equating to a density of ~ 1.80 g mL-1. Samples were then loaded into 4.9 mL OptiSeal tubes (Beckman Coulter, Brea, CA, USA), weighed to ensure proper balance, placed into a VTi 65.2 vertical rotor (Beckman Coulter), and spun at 37,000 r.p.m. at 20°C for 64 hr using an Optima L-80 XP ultracentrifuge (Beckman Coulter). Each gradient was fractionated into 12 tubes of approximately 410 μL each. The refractive index of each fraction was measured to determine density as described by Lueders, 2010 (25; Equation 1). RNA was precipitated with ice-cold isopropanol and the pellet was washed with ice-cold 70% ethanol as also described in (25). Following centrifugation, the RNA concentration of each fraction was determined using the RiboGreen quantification kit (Invitrogen) and a Gemini XPS plate reader (Molecular Devices, Sunnyvale, CA, USA). The RNA concentration was used as the guide for selecting which fractions to sequence. Quantitative reverse transcription polymerase chain reaction (qRT-PCR) was used to determine the quantity (copy number) of the 16S rRNA gene within each fraction using the KAPA Biosystems SYBR FAST One Step RT-qPCR (ABI Prism) and the Universal 16S rRNA primers, Pro341F and Pro805R primers (74). E. coli ribosomal RNA was used to construct a standard curve. Results were used to determine the ratio of maximum quantity of 16S rRNA within each set of fractions, that is the ratio of 16S rRNA in each fraction to the maximum copy number across all fractions (25, 29, 30; Figs S3-S5).

**Equation 1.** Lueders equation for fraction density (CsTFA + Gradient Buffer + Formamide)

𝑦 = −1893.9820𝑥^2^+5230.8018𝑥−3609.6804

**Metagenomic and Metatranscriptomic Library Construction**

Library construction for RNA-SIP fractions was carried out as described in Fortunato and Huber (2016; 8). Briefly, double stranded cDNA was constructed using SuperScript III First-strand synthesis system (Invitrogen, Grand Island, NY, USA) and mRNA second strand synthesis module (NEB, Ipswich, MA, USA). The synthesized double stranded cDNA was sheared to a fragment size of 175 bp using a Covaris S-series sonicator (Woburn, MA, USA). SIP metatranscriptomic library construction was completed using the Ovation Ultralow Library Complete Prokaryotic RNA-Seq DR multiplex system (Nugen, San Carlos, CA, USA) following manufacturer instructions, amplified using 15 cycles. Ribosomal RNA was not removed before construction of libraries. Sequencing was performed on an Illumina HiSeq 1000 at the W.M. Keck sequencing facility at the Marine Biological Laboratory in Woods Hole, MA. All libraries were paired-end, with a 30 bp overlap, resulting in an average merged read length of 110 bp.

The 47 mm flat filters that were collected and preserved in situ for metagenomic and metatranscriptomic sequencing were processed according to (10). The filters were first cut in half with a sterile razor, half used for DNA extraction for metagenomic sequencing and half used for RNA extraction for metatranscriptome sequencing. For DNA extraction, the DNA filter was rinsed with sterile PBS to remove the salts from the RNAlater and was then extracted using a phenol-chloroform method adapted from (75) and (76). DNA was then sheared to a fragment size of 175 bp (2013 and 2014) or 275 bp (Anemone, Marker 33, Marker 113 metagenomes from 2015, plume, and background seawater) using a Covaris S-series sonicator. Metagenomic library construction was completed using the Ovation Ultralow Library DR multiplex system (Nugen) following manufacturer instructions. RNA was extracted using the mirVana miRNA isolation kit (Ambion). Ribosomal RNA removal, cDNA synthesis, and metatranscriptomic library preparation was carried out using the Ovation Complete Prokaryotic RNA-Seq DR multiplex system (Nugen) following manufacturer instructions.

**Metagenomic and Metatranscriptomic Sequencing**

Metagenomic and metatranscriptomic sequencing was done on an Illumina HiSeq 1000 in 2013 and 2014 and an Illuma NextSeq 500 in 2015. All libraries were paired-end, with a 30 bp overlap, resulting in an average merged read length of 110 bp for libraries from 2013 and 2014 and 151 bp for libraries from 2015. Metagenomic and metatranscriptomic sequencing was carried out at the W.M. Keck sequencing facility at the Marine Biological Laboratory in Woods Hole, Massachusetts.

**Differential Expression Analysis**

*Defining contrasts for differential expression*

DESeq2 statistically compares abundances of genes across experimental conditions (34). A common implementation of DESeq2 is for the quantitative analysis of RNA-seq data to test the null hypothesis that gene expression is unchanged between two groups (34, 35, 37). However, DESeq2 can also robustly interpret experimental designs with multiple variables and/or high within-group variability in gene expression and imbalanced sample replication. We examined how temperature, time, and/or the vent’s unique chemistry affect expression patterns and metabolisms of chemoautotrophs. Since all of the data examined for differential expression are from ^13^C-incubated samples, it would be erroneous to consider the unmanipulated fluids as the control for each of these models because that does not allow us to disentangle the interaction of ^13^C-isotope spike + incubation temperature + incubation time as an influence on the expression profiles. We customized our input and models to reduce stochasticity in gene dispersion estimates, accommodate the three conditions (vent, temperature, and year) with multiple levels, and address specific hypotheses about chemoautotrophy in these vents.

For example, to explore differential gene expression in chemoautotrophs at 80°C due to vent chemistry (i.e. between Marker 113, Marker 33, and Anemone), we subset the data to reduce variability in modeled gene dispersion estimates as a result of temperature (i.e. remove the 30°C and 55°C profiles), and remove the interaction effect of time by considering as replicates the vent-specific samples across different years incubated at 80°C. In this examination, the baseline expected gene expression profile is calculated from the ^13^C-enriched samples incubated at 80°C (n=7).

When determining differential expression due to incubation temperature in each vent, we still aimed to be able to compare these results between vents. To enable these comparisons, we used all 21 of the ^13^C-enriched samples to estimate gene baseline expression. Then in the model, we removed year as a variable, subset the dataset by vent, and allowed for temperature to vary. These contrasts can then be built for each comparison of interest as described in the main manuscript (37).

*Sample ‘replicates’*

For the vent-specific across-temperature comparisons, we considered timepoints and different years as ‘replicates’ (collapsing ^13^C-enriched metatranscriptomes by time). For the across-vents, temperature-specific comparisons, we collapsed ^13^C-enriched metatranscriptomes by time again. For the within-vent, within-temperature, across years comparison, we considered timepoints of ^13^C-enriched metatranscriptomes as replicates.

*Choice of baseline*

As noted, a baseline of the unmanipulated metatranscriptomic expression was not used, because it would not accurately capture comparisons of an isotope-spiked sample across temperature and between years. Instead, a baseline (grouped by specific comparison) of averaged ^13^C-enriched transcript abundance for each condition was chosen (13H only), instead of an average of ^12^C-light control transcripts + ^13^C-enriched transcripts (13H + 12L) or the average of only ^12^C-light transcripts (12L only) from the RNA-SIP experiments. These normalized expression analyses support the determination to use only ^13^C-enriched samples in our DE contrasts, since the patterns (number of SDE genes and in which direction) in each comparison are not substantially changed when we choose a different contrast (13H vs 13H + 12L), and 13H contrasts produced a slightly higher number of SDE genes than the 13H + 12L baseline.
