## Supplementary material for "Metabolic and Population Profiles of Active Subseafloor Autotrophs in Young Oceanic Crust at Deep-Sea Hydrothermal Vents": Main Figures

**Table 1.** All RNA-SIP experiments, showing vent name, year, temperature of experiment, and time points collected.

| **Vent** | **Year** | **30 °C** | **55 °C** | **80 °C** |
| --- | --- | --- | --- | --- |
| **Marker 113** | 2013 | 36 hr | 36 hr | 18 hr* |
| **Marker 113** | 2014 | n.d. | 18 hr, 36 hr | 9 hr |
| **Anemone** | 2013 | n.d. | n.d. | 18 hr* |
| **Anemone** | 2014 | 24 hr, 36 hr | 18 hr, 36 hr | 18 hr |
| **Marker 33** | 2013 | 36 hr | 36 hr | 36 hr |
| **Marker 33** | 2014 | 24 hr, 36 hr | 18 hr, 36 hr | 9 hr, 18 hr |
| n.d. = no data; * = 1 L LVWS sample |  |  |  |  |

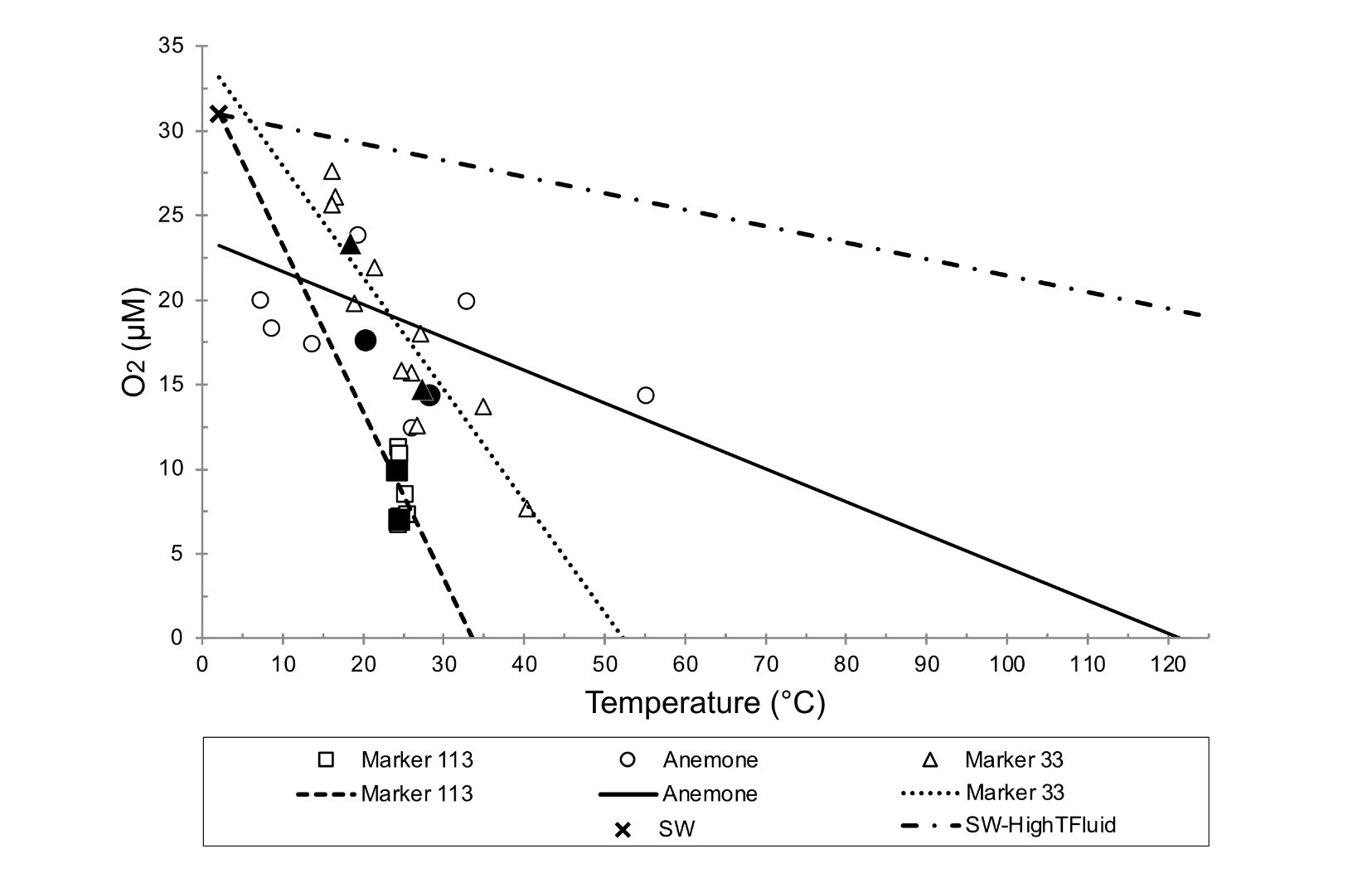

**Figure 1.** Figure 1. *In situ* oxygen versus temperature for diffuse vents sampled for this study. Near-bottom seawater averages 31 µmol/kg (+/- 2.2). Linear regression lines are drawn for each of the three vent sites for the two years sampled, including the ambient seawater point in the regression. The SW-HighTFluid line is between seawater and a hypothetical zero-oxygen end-member at 325°C. Solid data points indicate the exact fluid sampling bag used in the SIP experiment.

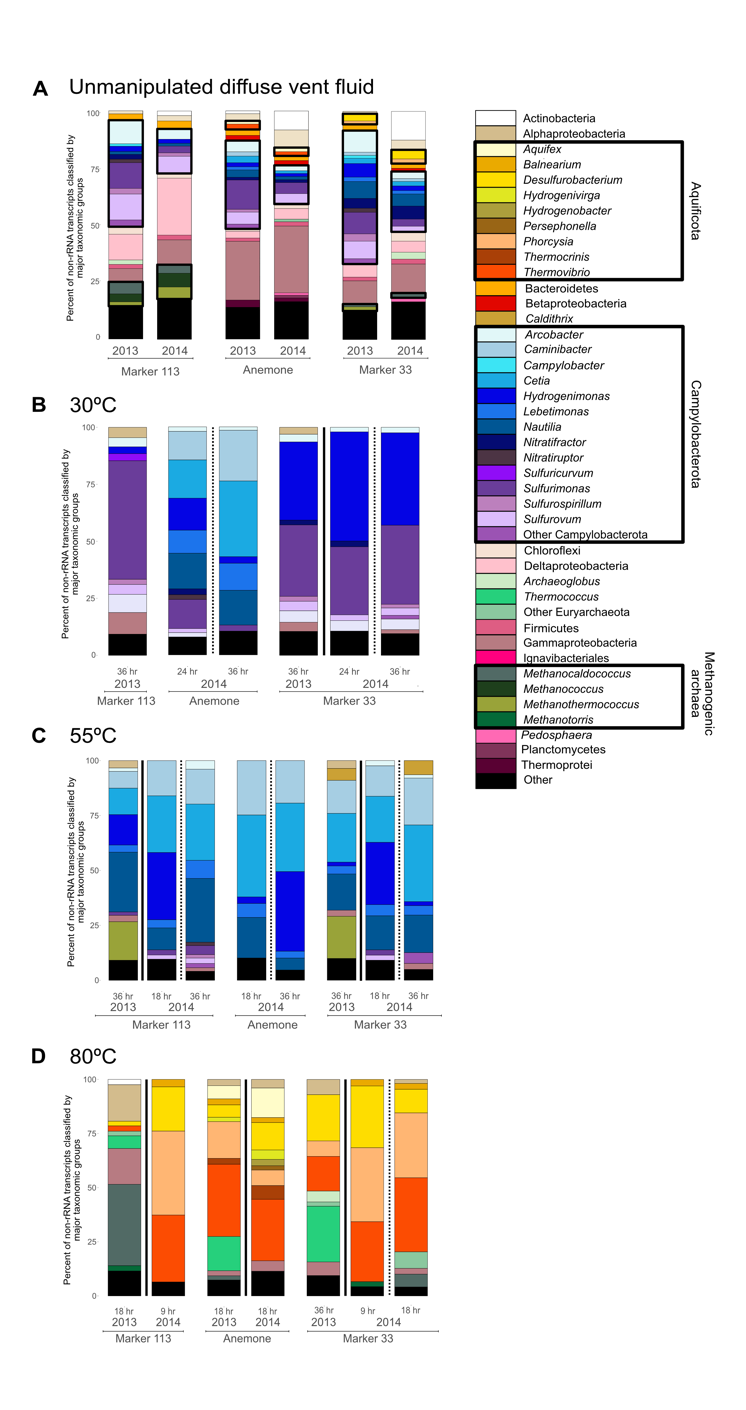

**Figure 2.** Relative abundance of major taxonomic groups of **A**. Annotated non-rRNA transcripts from unmanipulated fluid metatranscriptomes and annotated non-rRNA transcripts from the ^13^C-enriched fraction of RNA-SIP experiments at **B**. 30°C, **C**. 55°C, and **D**. 80°C. Black boxes indicate the main autotrophic taxonomic groups recovered from the ^13^C-enriched fraction of RNA-SIP metatranscriptomes and are drawn onto **3A** to show how these taxa make up a small proportion of the overall unmanipulated fluid metatranscriptome to then become the key autotrophs in the ^13^C-enriched fraction of RNA-SIP metatranscriptomes in **3B-D.**

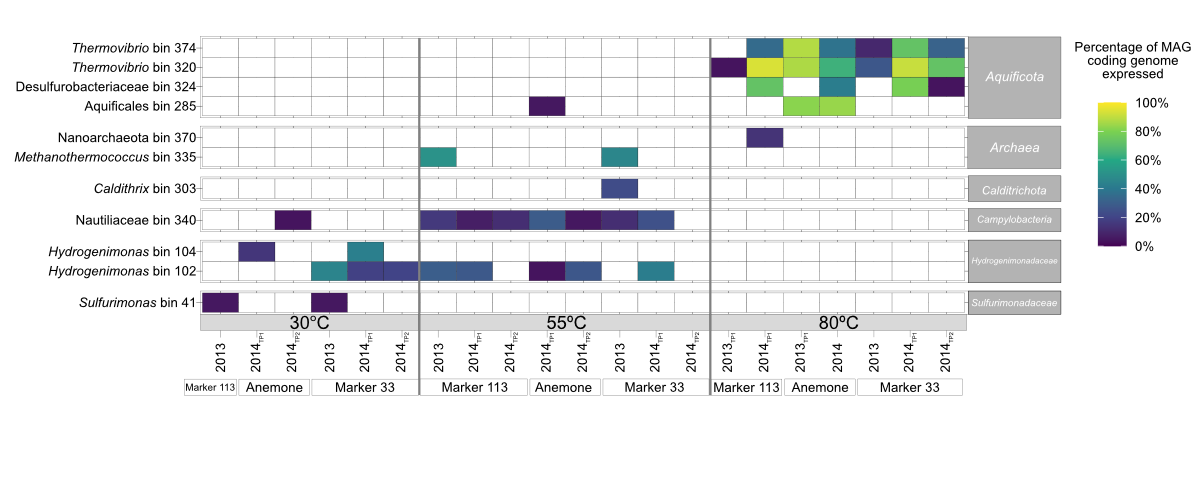

**Figure 3.** Metagenome Assembled Genomes (MAGs) selected based on having >4% of coding genome expressed in 30, 55 and 80°C ^13^C-enriched fraction of RNA-SIP experiments. Percent of coding genome expressed = length of expressed genes(bp)/genome length (bp).

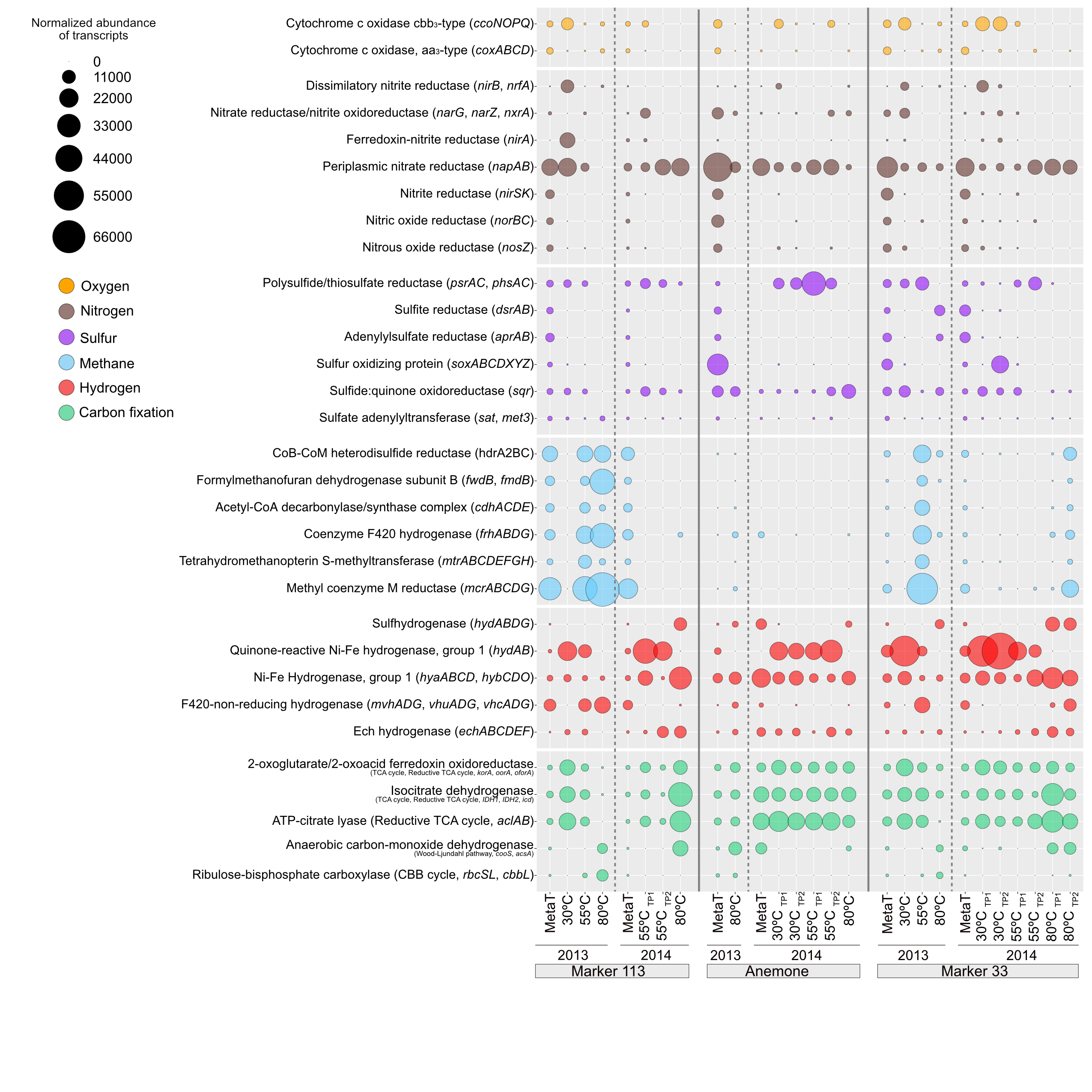

**Figure 4.** Normalized abundance of non-rRNA transcripts annotated in the unmanipulated diffuse fluid metatranscriptomes (“MetaT”) and ^13^C-enriched fractions of RNA-SIP metatranscriptomes for key marker genes involving oxygen, nitrogen, sulfur, methane, hydrogen, and carbon fixation. Bubble sizes represent the transcripts per million reads (TPM) attributed to each marker gene.

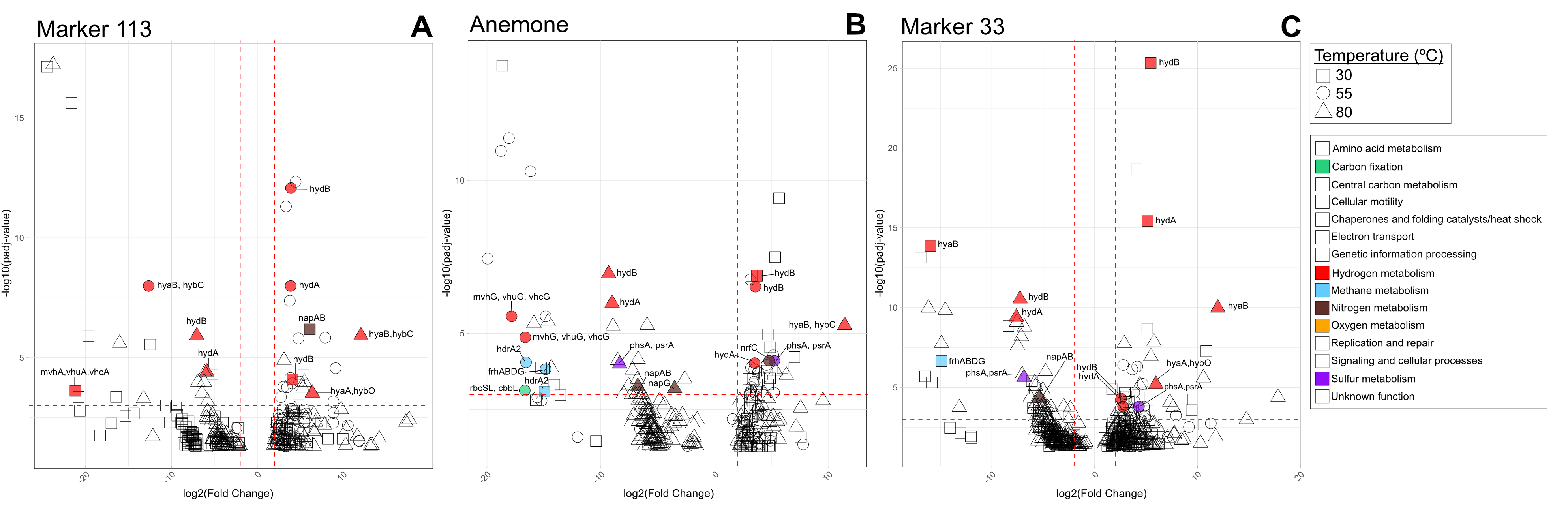
**Figure 5.** Volcano plots showing the log_2_FC (degree of differential expression) across experimental temperatures within each vent at **A.** Marker 113, **B.** Anemone, and **C.** Marker 33. Colored and labeled points correspond to genes involved in carbon fixation, hydrogen, methane, nitrogen, oxygen, and sulfur metabolism that had a padj < 10^-3^ and log_2_FC > |2| (padj; p-value adjusted for multiple tests using the Benjamini-Hochberg procedure to control the false discovery rate (FDR).
